# Exploring the Role of Kv1.3 and MAPK14 in Mediating Microglial Oxidative Stress and Neuroinflammation Following Organic Dust Exposure

**DOI:** 10.1101/2025.01.10.632439

**Authors:** Nyzil Massey, Sanjana Mahadev Bhat, Denusha Shrestha, Emir Malovic, Locke A. Karriker, Shivani Choudhary, Alan P. Robertson, Hai Minh Nguyen, Heike Wulff, Anumantha G. Kanthasamy, Chandrashekhar Charavaryamath

## Abstract

**Background:** Agricultural contaminants, including inhaled organic dust (OD) and gases, are known to cause inflammation in the lungs and the brain. We investigated the role of the potassium channel Kv1.3 in organic dust (OD)-induced neuroinflammation models. Kv1.3 channels play a multifaceted role in microglial immune modulation, cancer, neurodegenerative diseases, and constitute a potential therapeutic target.

**Methods:** We used *in vivo* (C57BL/6 mice), *in vitro* (microglial cell line, primary microglia), and *ex vivo* (brain slice culture) models of OD-induced neuroinflammation. A sterile OD extract (ODE) was prepared, and mice were exposed to either normal saline or ODE intra-nasally for 5 weeks (5 days/week) to simulate an occupational exposure scenario. Primary microglia were isolated from neonatal mice for total RNA sequencing (RNA-seq). The ODE-induced expression of Kv1.3 was quantified using *in vitro* and *ex vivo* models with and without PAP-1 treatments. Exposure-induced changes in cytokines and reactive species markers were measured. Using western blot, we quantified phosphorylated p38 MAPK14 (p-p38 MAPK) and NOX2. We measured the currents through Kv1.3 channels using a microglial patch-clamp assay.

**Results:** Exposure to ODE increased the expression of Kv1.3 and p-p38 MAPK in mouse microglia without affecting the Kv1.3 currents at the cell surface. Exposure increased the levels of inflammatory cytokines and NOX2. Kv1.3 inhibition with PAP-1 decreased inflammatory markers (TNF-α and IL-6), levels of Kv1.3, p-p38 MAPK, NOX2, and nitrites.

**Conclusion:** Our study revealed that pharmacological inhibition of Kv1.3 potassium channels reduces ODE-induced neuroinflammation by decreasing inflammatory and oxidative stress markers.

## Introduction

Approximately one-quarter of the global workforce is engaged in agriculture (Roser, 2024) and with 170,000 deaths per year, farm work is one of the most dangerous professions (International Labor Organization, 2024). The increasing global population and ever-increasing demand for more affordable protein sources have transformed the animal food production sector into efficient large-scale production units. These animal production facilities are known to produce and store many contaminants including organic dust (OD) and various irritant gases on site (Nordgren & Charavaryamath, 2018; Poole et al., 2024; Sethi et al., 2017). Inhaled OD is currently under investigation for its adverse effects, specifically its role in triggering respiratory and neurological inflammation (Massey et al., 2021a; Massey et al., 2022). This dust comprises a complex blend of gases such as methane, ammonia, and hydrogen sulfide (H_2_S), particulate matter (PM) ranging from 0.01 to 1000 µm in size, and numerous pathogen-associated molecular patterns (PAMPs) such as lipopolysaccharide (LPS) and peptidoglycan (Nordgren & Charavaryamath, 2018; Poole et al., 2024). The majority of the current research on the health impacts of OD is focused on the respiratory effects (Bhat et al., 2019; Bhat et al., 2022; Mahadev Bhat et al., 2021; Poole et al., 2024; Schwab et al., 2024; Shrestha et al., 2021; Shrestha et al., 2022), while our group has started investigating how inhaled OD exposure leads to neuroinflammation (Massey et al., 2019; Massey et al., 2021a; Massey et al., 2022). Despite these efforts, the cellular and molecular underpinnings of neuroinflammation due to OD remain largely elusive.

Research has established a link between farming activities in the United States and a heightened occurrence of neurodegenerative diseases, particularly in the Midwest (Wright Willis et al., 2010). Supporting this, research conducted in Iowa (Arora et al., 2021) suggests that individuals employed in agriculture are more likely to develop dementia, implicating occupational exposure in agriculture as a key factor in triggering neuroinflammation. Several studies have described how inhaled diesel exhaust particles (DEP), pesticides, and PM can negatively impact brain health (Levesque et al., 2011), (Sánchez-Santed et al., 2016), (Roux et al., 2017). However, how respiratory exposure to OD leads to neuroinflammation and subsequent cognitive impairments is not fully understood.

Our previous work has demonstrated that exposure of mouse brain microglial cells to OD extract (ODE) triggers a pro-inflammatory response via HMGB1-RAGE signaling (Massey et al., 2019). Either blocking nucleocytoplasmic translocation of HMGB1 via ethyl pyruvate or siRNA-mediated reduction in HMGB1 expression curtailed the pro-inflammatory response (Massey et al., 2019). Additionally, our use of mitoapocynin (MA), a mitochondria-targeted NOX2 inhibitor (Ghosh, Langley, et al. 2016) decreased the ODE-induced reactive nitrogen species (RNS) production (Massey et al., 2019). Our published work also highlights the role of mitochondrial dysfunction exposure of human THP1 cells to ODE (Mahadev Bhat et al., 2021). Recently, using both *in vitro* microglial cell lines and *ex vivo* brain slice cultures, we have shown that ODE exposure results in mitochondrial DNA (mtDNA) leakage into the cytosol, suggesting a mitochondrial membrane compromise (Massey et al., 2021b). This extramitochondrial mtDNA activates the cGAS-STING pathway, an innate immune mechanism for detecting foreign DNA (Wu et al., 2013). We have demonstrated that MA treatment and the siRNA-mediated reduction of STING mRNA greatly diminish the ODE-induced inflammatory response (Massey et al., 2021a).

Our *in vitro* and *ex vivo* models of OD exposure have uncovered mechanisms of exposure-induced cellular inflammation and ultra-structural damage to mouse microglia (Massey et al., 2019; Massey et al., 2021a; Massey et al., 2022). To provide translational relevance, we used an established mouse model that was originally designed by Dr. Poole’s group and later used by our group (Charavaryamath et al., 2005; Poole et al., 2009). Our results from the mouse experiments concluded that prolonged intranasal ODE exposure elicited an inflammatory response in the mouse brain, and treatment with oral MA provided a partial neuroprotection (Massey et al., 2022).

Since microglial cells are the chief innate immune cells in the brain, a comprehensive understanding of how they respond to ODE exposure is crucial. Hence, an RNA sequence analysis of microglia will likely provide the transcriptomic details of the exposure-induced neuroinflammation. The current study began with ODE exposure to identify gene expression changes in mouse brain microglia through RNA-seq analysis. This initial analysis revealed differential expression patterns, highlighting upregulated pathways associated with biological processes (BP) and molecular functions (MF). These findings provided the basis for our subsequent investigation into the specific roles of the Kv1.3 and MAPK14 pathways. Voltage-gated potassium (Kv) ion-channels have recently been identified as key players in neuroinflammatory conditions and other disorders (Yi-Je Chen et al., 2021; Izumi Maezawa et al., 2018; Nelson, 2006; Sarkar et al., 2020). The *kcna3,* gene that encodes for Kv1.3 is predominantly expressed in immune cells. Microglial cells express significant amounts of Kv1.3 channels (H. M. Nguyen et al., 2017) and play a central role in neuroinflammation and neurodegeneration (Castro-Gomez & Heneka, 2024; Y. J. Chen et al., 2021; Chen et al., 2018; Krause & Müller, 2010; I. Maezawa et al., 2018; Hai M. Nguyen et al., 2017).

We have previously shown that microglia, upon exposure to ODE, generate reactive oxygen species (ROS) and nitrite (Massey et al., 2019). The interaction between superoxide and nitrite results in the formation of peroxynitrite, a compound with neurotoxic properties. As a potent oxidizing and nitrating substance, peroxynitrite is implicated in the advancement of neurodegenerative diseases, primarily through its ability to induce neuronal death via the Kv1.3 channels (Fordyce et al., 2005; Torreilles et al., 1999).

The role of Kv1.3 in LPS-induced microglial neuroinflammation has been established (Di Lucente et al., 2018; Fordyce et al., 2005). Since OD is a complex mixture containing LPS (Nordgren & Charavaryamath, 2018) and other toxicants, it can trigger a multifaceted signaling pathway. The p38-α subtype of mitogen-activated protein kinases (MAPK14) detects cellular damage, and leads to the production of cytokines and RNS through the NF-κB pathway (Ashraf et al., 2014; Begoña Canovas & Angel R. Nebreda, 2021; Fordyce et al., 2005). The phosphorylated form of MAPK14 (pMAPK14) plays a crucial role in microglial neurotoxic signaling (Fordyce et al., 2005). pMAPK14 influences the expression of Kv1.3 channels during HIV-1 glycoprotein 120 infections (J. Liu et al., 2012). Moreover, PAP-1 (5-(4-phenoxybutoxy)psoralen) selectively inhibits Kv1.3 channels in a state-dependent manner without showing cytotoxic or phototoxic effects. This compound has been effectively utilized to block Kv1.3 channels in a Parkinson’s disease model, thereby reducing neuroinflammation and neurodegeneration (Sarkar et al., 2020). Based on these premises, we hypothesize that OD exposure increases the expression of Kv1.3 channels and pMAPK14, leading to increased production of reactive oxygen species (ROS), nitrite, and inflammatory cytokines.

First, we used a mouse model of intra-nasal ODE exposure and performed RNA sequence analysis of brain microglia. Our results confirmed that, ODE exposure upregulates Kv1.3 expression. Next, we tested our hypothesis using an *in vitro* microglial cell line, primary microglial cultures, and an *ex vivo* brain slice culture (BSC) model. To clarify the role of Kv1.3 in OD-induced neuroinflammation, we utilized PAP-1 as a pharmacological inhibitor. We concluded that, PAP-1 mediated targeting of Kv1.3 channels reduces the ODE-induced neuroinflammation by decreasing inflammatory and oxidative stress markers.

## Methods and materials

### Preparation of organic dust extract

All experiments were conducted in accordance with an approved protocol from the Institutional Biosafety Committee (IBC protocol# 19-004) of Iowa State University. Settled swine barn dust (representing OD) was collected from various swine production units into sealed bags with a desiccant and transported on ice to the laboratory. Organic dust extract (ODE) was prepared as per a published protocol (Romberger et al., 2002). Briefly, dust samples were weighed, and for every gram of dust, 10 mL HBSS, no calcium, no magnesium (Thermo Fisher, cat. # 14170112) was added, stirred, and allowed to stand at room temperature for 60 min. The mixture was centrifuged (1365 g, 4° C) for 20 min, supernatant collected, and the pellet was discarded. The supernatant was centrifuged again with the same conditions, the pellet discarded and recovered supernatant was filtered using a 0.22 µm filter and stored at −80° C until used. This stock was considered 100% and diluted in cell culture medium to prepare a 1% v/v solution to use in our *in vitro* experiments (Table 1). We routinely quantify the LPS content of our ODE samples using Pyrochrome® Kinetic Chromogenic Endotoxin Assay (Cape Cod, cat. # CG-1500-5) as per the instructions. We have previously published the LPS content of several of our ODE samples (Bhat et al., 2019).

**Table 1.** Treatments.

| Treatment group | *in vivo* | *ex vivo* | *in vitro* |
| --- | --- | --- | --- |
| Control | Normal saline<br>(Intranasal) | BSCs Media | BV2 & primary<br>microglia Media |
| ODE | 12.5 % (5-day x 5<br>weeks)<br>(Intranasal) | 1% v/v (5 day) | 1% v/v (48 h) |
| ODE+PAP-1 | Not done | 1μM PAP-1 (Co-treatment) | 1μM PAP-1 (Co-treatment) |
| Lipopolysaccharide<br>(positive control) | Not done | Not done | 100ng/mL |

### Chemicals and reagents for *in-vivo* and *ex-vivo* culture

Dulbecco’s Minimum Essential Medium (DMEM) (Thermo Fisher, cat. # 11965092), Penicillin-Streptomycin (10,000 U/mL) (Thermo Fisher, cat. # 15140122), L-Glutamine (200 mM) (Thermo Fisher, cat. # 25030081), and Trypsin-EDTA (0.25%), phenol red (Thermo Fisher, cat. # 25200072), and Fetal Bovine Serum (FBS) (Atlanta Biologicals, Flowery Branch, GA, cat. # S11150H, Lot # A17002) were utilized for *in-vivo* and *ex-vivo* culture. PAP-1 [5-(4-phenoxybutoxy) psoralen] was synthesized in Dr. Wulff’s laboratory as described (Schmitz et al., 2005) and made into a 10 mM/L stock solution in DMSO and stored at −20° C. PAP-1 was used as one of the co-treatments with ODE (1% v/v) (Table 1).

### Animal care, treatment, and euthanasia

Mouse exposure experiments were performed as per approved protocol (IACUC-19-250) by the Institutional Animal Care and Use Committee (IACUC) at Iowa State University (Ames, IA, USA). Eight-week-old male C57BL/6 mice (3 mice/group), obtained from Charles River, were housed under the following standard conditions: constant temperature (22 ± 1°C), humidity (relative, 30%), and a 12-h light/dark cycle. The animals were assigned randomly by a coin flip to the treatment groups using the mean weight of the group as the criteria. The mean weight of each group was not significantly different from other groups. Investigators involved in data collection and analysis were blinded to the treatment groups. After acclimatization for 7 days (week 0), mice were either intranasally administered with normal saline (0.9% w/v) or 12.5 % ODE (25 µL into each nostril, total of 50 µL/mouse/day) 5 days/week (Monday-Friday) for a total of 5 weeks (Table 1). Mice were euthanized following 5 weeks of intra-nasal exposure, using a pressurized CO_2_ chamber (AVMA-approved method), and freshly dissected brains were immediately harvested in ice-cold PBS. All the treatments were performed as per our previously published work (Massey et al., 2022).

### Preparation of microglial single-cell suspension and myelin removal

Meninges were removed from mice brain and the brains were cut into small pieces (<1 mm) with a scalpel and then placed in ice-cold Dulbecco’s modified Eagle’s medium/F-12 nutrient mixture DMEM/F-12 (Thermo Fisher, cat. # 3625) supplemented with 10% heat-inactivated fetal bovine serum (FBS), 50 U/mL penicillin, 50 μg/mL streptomycin, 2 mM L-glutamine, 100 μM non-essential amino acids, and Sodium Pyruvate (100 mM) (Thermo Fisher, cat. # 11360070) also referred to as sample preparation medium. The tissue pieces were incubated after adding 0.25% trypsin-EDTA in a 37°C water bath for 30 min with gentle agitation. A single-cell suspension of the digested brain tissue was prepared by gentle trituration and passing through 70 μm reversible Strainers (STEMCELL Technologies, cat. #27260) to remove tissue debris and aggregates. Single-cell suspension was centrifuged at 300 x g for 10 minutes at room temperature or at 2-8°C, with the brake on low. The supernatant was carefully removed and discarded. 30% Percoll® (GE Healthcare, cat.#. 17-0891-01) solution was added to the pellet and centrifuged at 700 x g for 10 minutes at room temperature or 2-8°C. The upper myelin layer was carefully removed and discarded. The pellet was washed with sample preparation medium once and resuspended to the final concentration of 2.5 x 10^7^ cells/mL.

### Column-free microglial isolation from single-cell suspension

Microglia were isolated from freshly dissected mice brain via EasySep mouse CD11b positive selection kit II (STEMCELL Technologies, cat. #. 18970). The single cell suspension was resuspended at a density of up to 2.5 x 10^7^ cells/mL in the separation medium (PBS containing 2% FBS and 1 mM EDTA, with no calcium or magnesium) and transferred to a 5 mL polystyrene round-bottom tube (Corning, cat. # 352058). The microglia were then isolated as per the manufacturer’s protocol. Anti-CD16/32; FcR blocker (BioLegends, cat. # 101301) was used to enhance the purity of the single cell suspension by preventing non-specific labeling of other cell types (Gordon et al., 2011).

### Mammalian cell culture and treatments (BV2 microglia and primary microglia)

BV2 microglial cell line derived from wild-type C57BL/6 mice (Halle, Hornung et al. 2008) was a kind gift from Dr. DT Golenbock (University of Massachusetts Medical School, Worcester, MA) to Dr. AG Kanthasamy. Microglial cells (BV2 microglia and primary microglia) were grown in T-75 flasks (1 x 10^6^ cells/flask), 12-well tissue culture plates coated with poly-D Lysine, and 24-well tissue culture plates (50 x 10^3^ cells/well). Cells were maintained in DMEM supplemented with 10% heat-inactivated fetal bovine serum (FBS), 50 U/mL penicillin, 50 µg/mL streptomycin, and 2 mM L-glutamine. Cells were incubated overnight before treatment. All the treatment group details are outlined in Table 1. Control group samples were collected at 0 h because the control group samples from 6, 24, and 48 h time points showed no differences in our pilot studies (Massey et al., 2019). All treatments were done for 48 hours as per previous publications (Massey et al., 2019; Massey et al., 2021a). LPS was used as a positive control for microglial stimulation.

### RNA sequencing data quantification and analysis of isolated microglia

Total RNA was extracted from isolated microglia using TRIzol reagent, and quality was assessed using an Agilent 2100 Bioanalyzer (Agilent Technologies, Santa Clara, CA, USA). RNA sequencing was performed at Iowa State University’s DNA Facility following library preparation. Raw reads in fastq.gz format from each sample were processed using Galaxy, an open-source platform for bioinformatics workflows (Galaxy Community, 2024). The initial step involved obtaining clean, high-quality reads by removing adapter sequences and filtering low-quality reads. This was performed using Trim Galore! (Galaxy version 0.6.10). Quality metrics such as Q20, Q30, and GC content were assessed using FastQC (Galaxy version 0.73+galaxy0) and MultiQC (Galaxy version 1.17+galaxy0) for clean reads. The high-quality reads were aligned to the mouse reference genome (e.g., GRCm39) using HISAT2 (Galaxy version 2.2.1+galaxy0). The genome reference files (FASTA and GTF annotation) were downloaded from Gencode and uploaded to Galaxy. Single-end alignment was performed with default settings, generating BAM files for each sample. BAM files contained chromosomal coordinates for the mapped reads. Mapped reads were assigned to genes using featureCounts (Galaxy version 2.0.3+galaxy1), a tool from the Subread package. It utilized built-in annotations to calculate read counts based on genomic start and end positions of each exon. Individual count files were generated for each sample, and a count matrix was created using Concatenate datasets (Galaxy version 1.0.0) to consolidate counts into a single table (genes in rows, samples in columns). The count matrix, gene annotation file (containing gene symbols and descriptions), and factor data file (e.g., Control vs. ODE) were input into the limma-voom pipeline implemented via the limma-voom (Galaxy version 3.50.0+galaxy0) tool. The voom function applies precision weights to the raw counts data, and edgeR normalization was used to calculate counts-per-million (CPM) values (Robinson et al., 2010). Genes with an adjusted p-value < 0.05 (corrected for multiple testing using the false discovery rate, FDR) were identified as differentially expressed genes (DEGs).

### DAVID enrichment and gene ontology (GO) analyses of DEGs

Lists of differentially expressed genes (DEGs) and their corresponding ENTREZ IDs (p ≤ 0.05) for each treatment group were uploaded to the DAVID (Database for Annotation, Visualization, and Integrated Discovery) tool. The species background was set to “Mus musculus”. Gene Ontology (GO) enrichment analyses were conducted to identify significantly enriched terms within the categories of biological processes (BP), molecular functions (MF), and cellular components (CC). The DAVID pathway viewer was utilized to explore and visualize these enriched pathways, providing functional annotation and insights into the biological significance of the DEGs.

### Brain slice cultures (BSCs)

The research outlined in this document adhered to the protocols sanctioned by the Institutional Animal Care and Use Committee of Iowa State University (IACUC protocol #18-290). The methodology for preparing mouse organotypic brain slices was in line with established procedures (Kondru et al., 2017). Following an approved animal breeding protocol (IACUC protocol #18-227), breeding pairs of C57BL/6 mice were obtained from The Jackson Laboratories, Bar Harbor, ME, and mating was initiated when the mice were approximately four weeks old. Offspring were nurtured by their parents until they reached the age of 9-12 days. At this stage, both male and female pups were humanely euthanized via cervical dislocation, a method endorsed by the AVMA for animals within this age range. The brain slices were then meticulously crafted from the freshly harvested brain matter using a Compresstome™ VF-300 microtome (Precisionary Instruments). This process involved positioning the entire brain along the mid-sagittal plane inside the Compresstome’s tube, previously filled with 2% agarose gel. The gel was rapidly set by cooling with a chilling block. Subsequently, the tube was placed into a slicing chamber containing a cold solution of Gey’s balanced salt solution enhanced with kynurenic acid (GBSSK) to prevent excitotoxicity. The GBSS was concocted by sequentially mixing components from concentrated stocks to achieve the desired final concentrations per liter: NaCl (8g), KCl (0.37g), Na_2_HPO_4_ (0.12g), CaCl_2_·2H_2_O (0.22g), KH_2_PO_4_ (0.09 g), MgSO_4_·7H_2_O (0.07g), MgCl_2_·6H_2_O (0.210 g), and NaHCO_3_ (0.227g). The Compresstome’s compression lip, situated in the cutting area, ensured the brain remained stable during the slicing process, which yielded 350-μm thick sections at a medium vibration setting. These BSCs were collected from the tube’s exit and relocated to a new plate containing fresh GBSSK. After their preparation, the brain slice cultures (BSCs) underwent a double wash in 6 mL of chilled GBSSK. They were then placed into 6-well plate inserts (Falcon, catalog #353090), each containing 3 to 4 slices, and incubated at 37 °C in a 5% CO_2_ humidified environment for a fortnight. The process of slicing brain tissue can damage neuronal and glial cells, leading to immediate gliosis post-preparation. To mitigate this problem, a pre-treatment incubation period for the BSCs is advised. A two-week incubation has been shown to alleviate sectioning-induced trauma (Croft & Noble, 2018; Kondru et al., 2017). During this period, the culture medium was refreshed bi-daily. Following the incubation, a five-day treatment was administered to the BSCs (Table 1).

### Western blot

Fresh brain tissue samples were collected from animals after dissection and cut into small pieces with a scalpel and placed in RIPA buffer. Tissues were further triturated into a suspension and sonicated to get a whole cell lysate. Total protein was estimated using Bradford assay. Equal amounts of proteins (20 μg/well) were resolved on 10% SDS-PAGE gels (Bio-Rad). Next, proteins were transferred to a nitrocellulose membrane, and the nonspecific binding sites were blocked for an hour with a blocking buffer specially formulated for fluorescent western blotting (Rockland Immunochemicals, Pottstown, PA). Membranes were incubated overnight at 4 °C with the respective primary antibodies (listed in Table 2). Next, membranes were incubated with the respective secondary donkey anti-rabbit IgG highly cross-adsorbed (A10043) or anti-mouse 680 Alexa Fluor antibodies (A21058, Thermo Fisher Scientific). Membranes were washed three times with PBS containing 0.05% Tween-20 and visualized on the Odyssey infrared imaging system. Band densities were normalized using β-actin (1:5000, Abcam; ab6276 or ab8227) as a loading control, and densitometry was performed (ImageJ, NIH).

**Table 2.** Antibodies used in Western blotting.

| Antibody | Catalog | Vendor |
| --- | --- | --- |
| KCNA3 | CSB-PA003118 | CUSABIO |
| pMAPK14 | CSB-PA000709 | CUSABIO |

### qPCR analysis

Total RNA from mice brains, BSCs, and cell culture samples was isolated using TRIzol^TM^ (Invitrogen, cat. # 15596-026) as per the manufacturer’s guidelines. RNA concentration was measured using NanoDrop, and the A260/A280 ratio was used to determine the RNA quality. Samples with an A260/A280 ratio (1.8-2.1) were considered acceptable and used for further analysis. 1 µg of RNA was reverse transcribed into cDNA using the superscript IV VILO Kit (Thermo Fisher Scientific, cat. # 11766050) following the manufacturer’s protocol. For qPCR, 5 µL of PowerUpTM SYBR Green Master mix (Thermo Fisher Scientific, cat. # 25742), 0.5 µL each of forward and reverse primers (10 µM), 3 µL of water and 1 µL of cDNA (1-10 ng) were used. The genes and their respective primer sequences used for qPCR analysis are listed in Table 3 (DEGs validation; *in vivo*) and Table 4 (Gene expression; *in vitro* and *ex vivo*). All primers were synthesized at Iowa State University’s DNA Facility. β-actin was used as a housekeeping gene. No template controls and dissociation curves were run for all experiments to exclude cross-contamination. CT values of the gene products of interest were normalized to housekeeping gene product CT values. Comparisons were made between experimental groups using the ΔCT method. Briefly, the ΔCT value was calculated for each sample (CT gene of interest minus CT β-actin). Then the calibrator value was averaged (ΔCT) for the control samples. The calibrator was subtracted from the ΔCT for each control and from the experimental sample to derive the ΔΔCT. The fold change was calculated as 2^ΔΔ^Ct (Livak & Schmittgen, 2001). The average fold change was calculated for each experimental group and for the full names of each gene, refer to supplementary data (Supplementary Table 3).

**Table 3.**
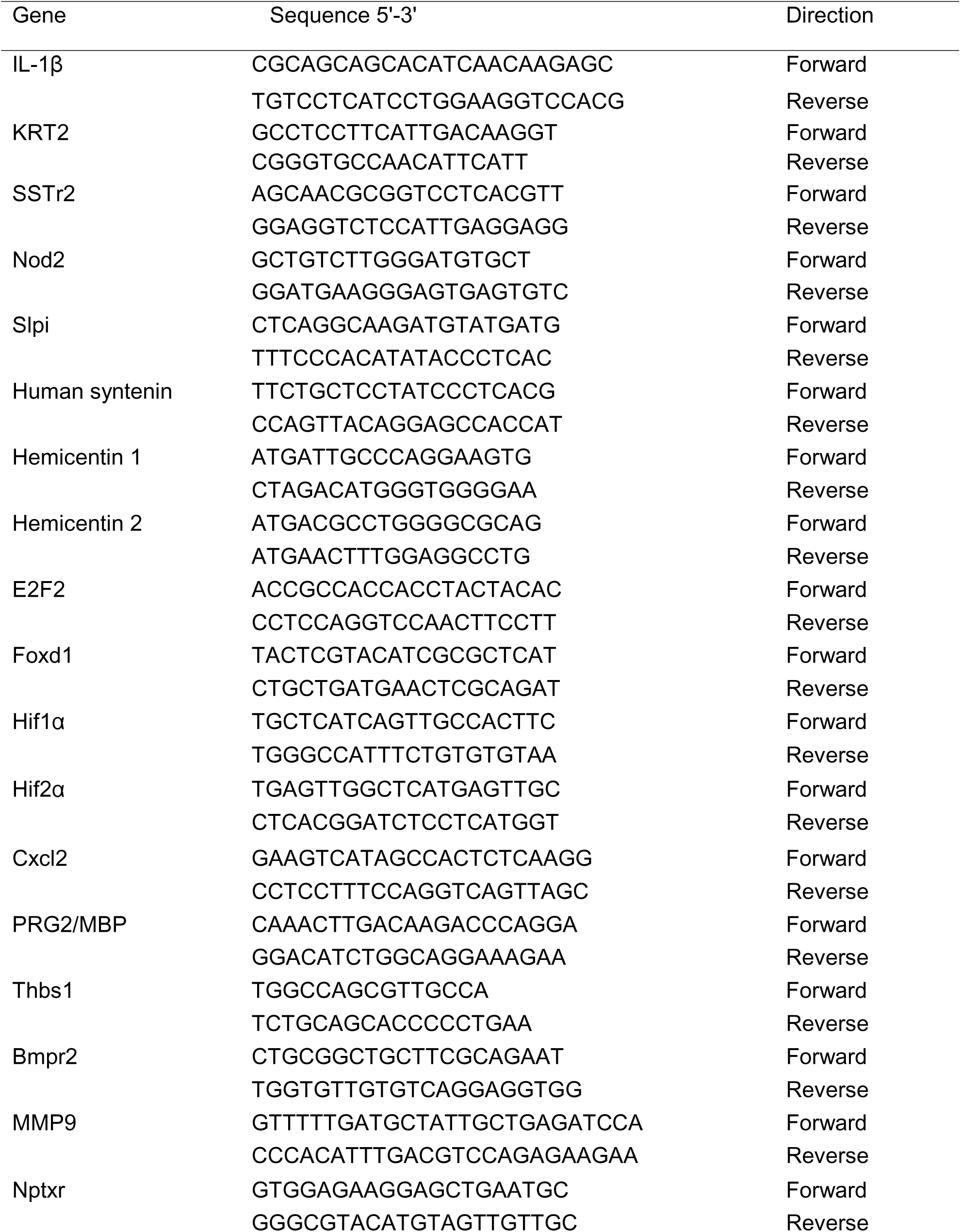
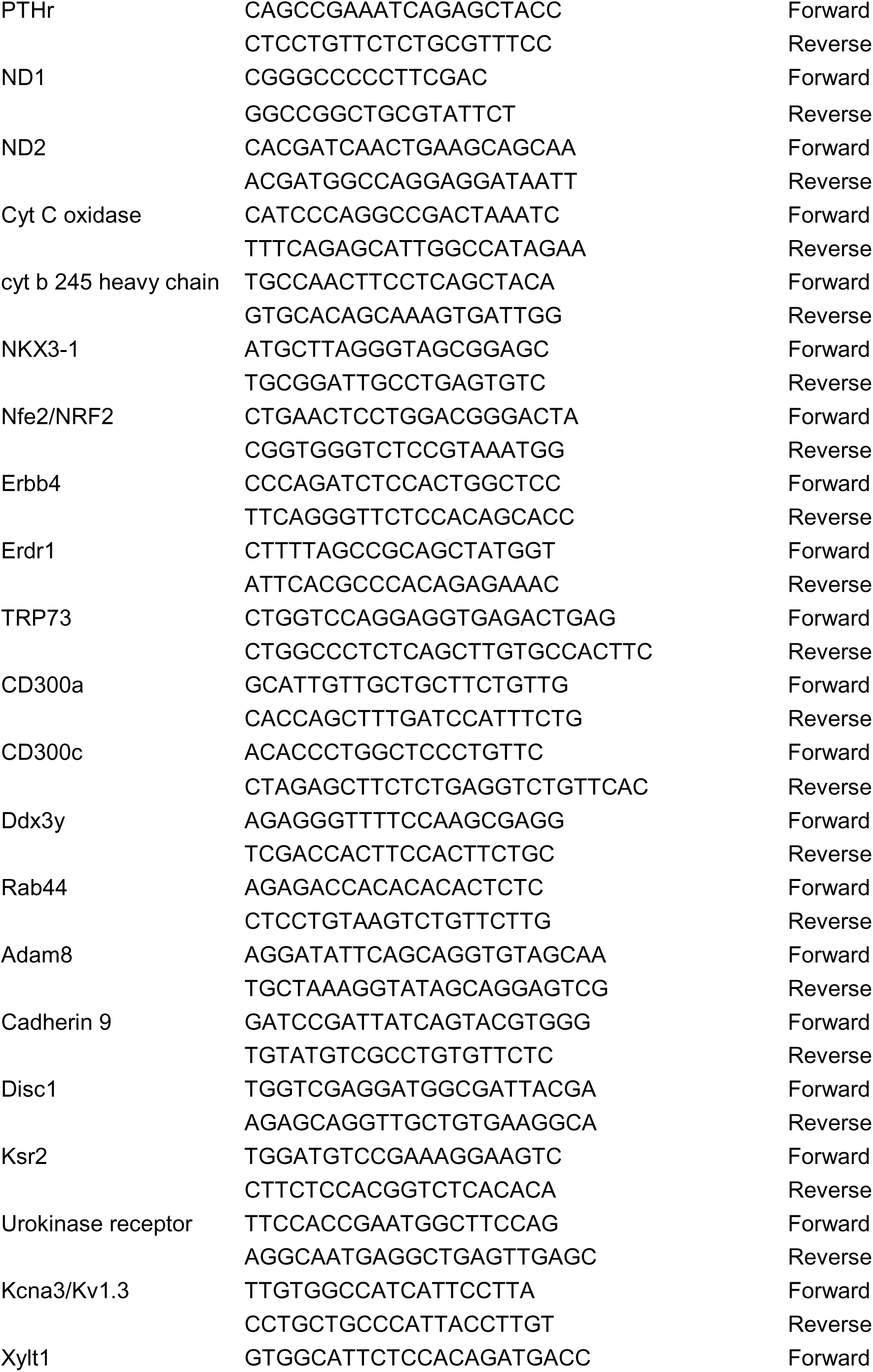
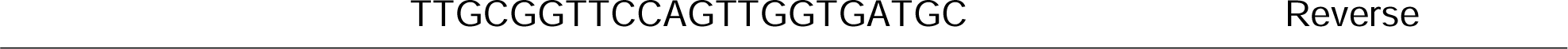
Primer sequences (*in vivo* for qPCR validation)

**Table 4.**
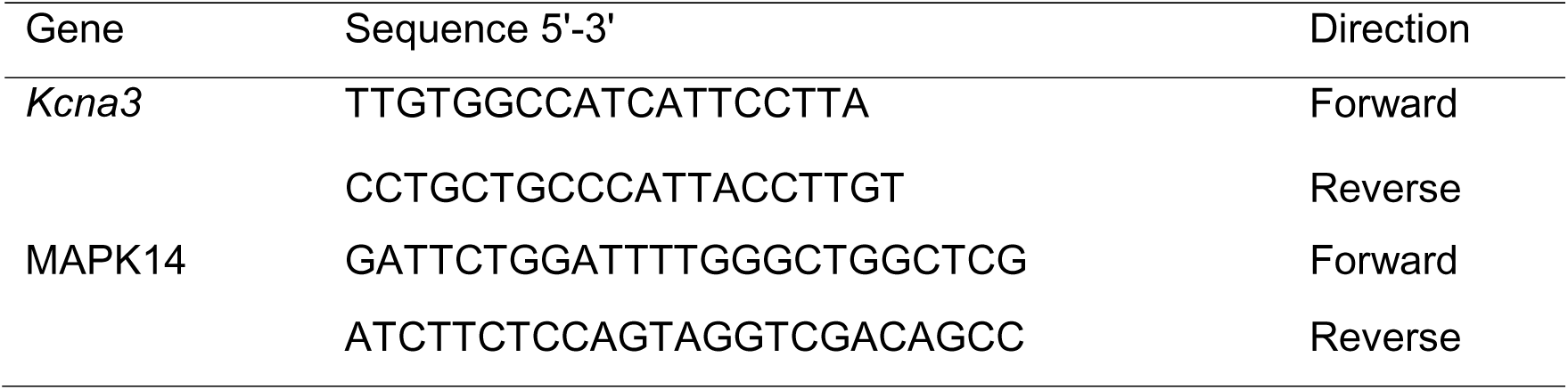
Primer sequences (in vitro and ex vivo)

### Cytokine analysis by ELISA

Culture media was collected following treatment of BSCs (5 days) and analyzed for TNF-α (Thermo Fisher Scientific, catalog # 88-7324-88) and IL-6 levels (Thermo Fisher Scientific, cat. # 88-7064-77) using ELISA kits by following recommended manufacturer’s protocols.

### Luminex assays for cytokine detection in BV2 microglia cell line and primary microglia

TNF-α, IL-6, IL-1β, IL-10, IL-17A, and IFNγ levels were assessed in supernatant culture media via Luminex assays. 40 μL of neat media was added to 40 μL of primary antibody conjugated to magnetic microspheres and incubated overnight at 4 °C in a clear-bottom, black 96-well plate. After incubation, each well was washed (3x) using a magnetic washer and then incubated for 1 h with secondary antibodies. Finally, samples were incubated for 30 minutes with streptavidin/phycoerythrin. A Bio-Plex reader was used to read the 96-well plates at 485 nm for excitation and 520 nm for emission. A standard curve of all the cytokines was prepared using standard cytokines.

### Reactive nitrogen species detection in the supernatants

We measured nitrite production (RNS marker) in BV2 microglia and BSCs using Griess reagent (Sigma-Aldrich). Briefly, the supernatant was collected following treatments (Table 1). Griess reagent was added to 50 mL of supernatant samples in a 98-well plate (1:1) and incubated at room temperature for 10 min until color development. A standard nitrite curve was prepared from sodium nitrite. The color change was analyzed by measuring absorbance at 540 nm using a microplate reader (Spectramax M2, Molecular devices). The absorbance values were expressed in µM, and data was analyzed using GraphPad Prism software.

### Immunohistochemical (IHC) analysis in BSCs

BSCs were treated with ODE or controls for 5 days as per our published study (Massey et al., 2021a). All the treatment groups are listed in Table 1., BSCs on inserts were carefully excised using a scalpel and transferred to new 12-well inserts facing upward. The BSCs were washed twice with PBS, fixed in 4% paraformaldehyde at room temperature for 30 minutes, and then incubated with ice-cold 20% methanol in PBS for an additional 5 minutes. Permeabilization was performed using 1% Triton X-100 in PBS at 4 °C for 12–18 hours. To minimize non-specific staining, BSCs were blocked with 20% BSA containing 0.1% Triton X-100 in PBS for 2–3 hours. The cells were then incubated overnight at 4 °C with primary antibodies (Table 5). After three washes with 5% BSA in PBS, BSCs were incubated with secondary antibodies for 12 hours at 4 °C. Following secondary antibody incubation, the cells were mounted using VECTASHIELD antifade mounting medium containing DAPI (Vector Labs, Burlingame, California) and covered with a cover glass. The cover glasses were sealed with nail polish, and the slides were imaged using a Nikon Eclipse TE2000-U microscope. Images were captured with a Photometrics Cool Snap CF camera and processed with HCImage software (Tucson, AZ).

**Table 5.**
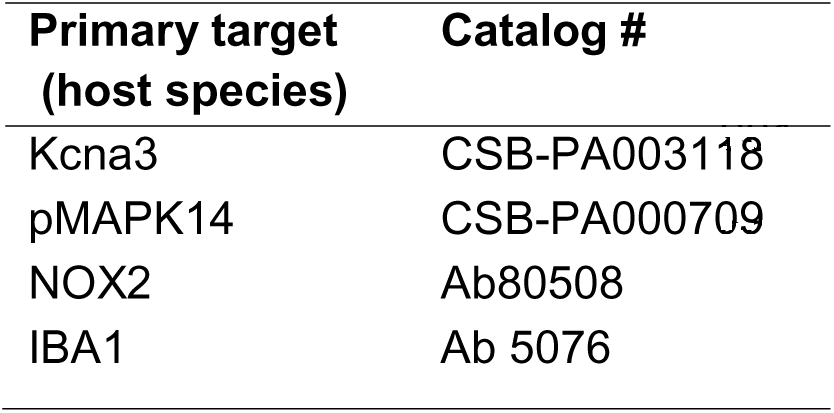
Antibodies used for immunostaining.

### Immunocytochemical (ICC) analysis in BV2 microglia and primary microglia

BV2 microglia and primary microglia were seeded into three 35-mm-diameter glass coverslips at the bottom of 24 well cell culture plates and maintained in DMEM (Thermo Fisher Scientific, Waltham, MA) supplemented with 10% heat-inactivated FBS, 50 U/mL of penicillin, 50 μg/mL of streptomycin and 2 mM L-glutamine and incubated overnight. Cells were treated as outlined in Table 1. After washing with ice-cold DPBS, blocking was performed with 10 % normal donkey serum (EMD Millipore, Burlington, Massachusetts) and 0.2% Triton-X100 in PBS for 1 hour at room temperature. Primary antibodies (Table 5) were prepared in antibody diluent solution (2.5% normal donkey serum, 0.25% sodium azide, 0.2% trition X 100 and PBS, Abcam, Cambridge, Massachusetts) and wells were incubated with primary antibodies overnight at 4^0^C. Next, each well was washed with PBS five times and then incubated with the corresponding secondary antibody for 1 h followed by mounting with VECTASHIELD antifade mounting medium containing 4’, 6-Diamidino-2-Phenylindole, Dihydrochloride (DAPI, Vector Labs, Burlingame, California) on glass sides. Slides were dried overnight and then imaged using a Nikon Eclipse C1 microscope.

### IHC and ICC Quantification

Five fields per coverslip were chosen randomly, and total stained cells were counted (ImageJ). Total staining intensity per field (cy3 or FITC) was measured using HC Imagesoftware (Hamamatsu Corp, Sewickley, Pennsylvania). The mean cell intensity was measured by dividing total intensity by the number of cells per field (Massey et al., 2019). The average mean intensity for each coverslip was calculated and plotted.

### Whole-cell patch clamping

Primary microglia were plated at 100,000–150,000 cells/well in 24-well plates for 8–12 hours before stimulation with either 300 ng/mL LPS or 1% ODE. After 24 or 48-hour, cells were detached by trypsinization, washed, attached to poly D lysine-coated glass coverslips, and then studied within 20–90 minutes after plating in the whole-cell mode of the patch-clamp technique with an EPC-10 HEKA amplifier (HEKA Elektronik). All currents were recorded in normal Ringer’s solution containing 160 mM NaCl, 4.5 mM KCl, 2 mM CaCl_2_, 1 mM MgCl_2_, and 10 mM HEPES (adjusted to pH 7.4 and 290–310 mOsm). Patch pipettes were pulled from soda-lime glass (micro-hematocrit tubes, Kimble Chase) to resistances of 2–3 MΩ when submerged in the bath solution and filled with a KF-based Ca^2+^-free internal pipette solution containing 145 mM KF, 2 mM MgCl_2_, 10 mM HEPES, and 10 mM EGTA (pH 7.2, 290–310 mOsm). Recording conditions were set up to isolate Kv1.3 currents. The use of KF avoids “contaminating” Kv currents with calcium-activated K^+^ currents or chloride currents. Currents were elicited with a “use-dependence protocol,” involving a train of ten 200-ms voltage steps from –80 to 40 mV applied at a frequency of 1 Hz, which identifies Kv1.3 by its characteristic use dependence (e.g., current amplitude declines rapidly when pulsed faster than the channels can recover from inactivation). In 25% of the cells, we then further pharmacologically confirmed that the current was predominantly carried by Kv1.3 by testing its sensitivity to PAP-1. Cell capacitance, which directly measures the cell-surface area, and access resistance were continuously monitored during recordings. Kv1.3 current density was calculated as the use-dependent current amplitude at 40 mV divided by the cell capacitance measured for individual cells (H. M. Nguyen et al., 2017).

### Statistical analysis

Data were expressed as mean ± SEM and analyzed by one-way ANOVA or two-way ANOVA followed by Tukey’s post hoc comparison tests (GraphPad Prism 10.0, La Jolla, California). A p-value of ≤ 0.05 was considered statistically significant. An asterisk (*) indicates a significant difference between controls and ODE-treated cells, whereas hashtag (#) indicates a significant difference between control and PAP-1 treatment. The *p-values* corresponding to asterisk/s or hashtag/s are listed in Table 6. The details of the statistical analysis applied for each experiment are listed (Supplementary Table 4).

**Table 6.**
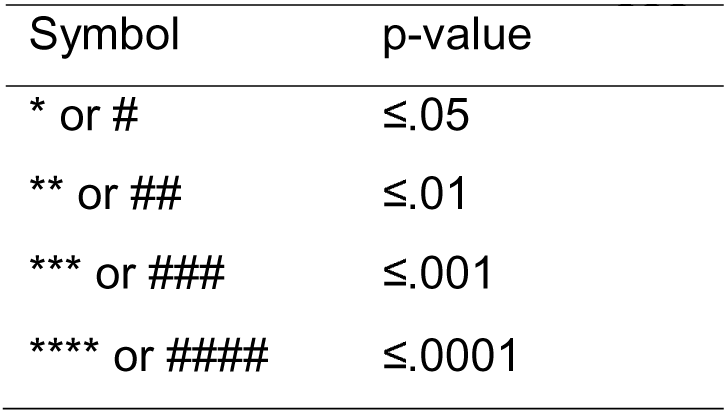
Symbols (asterisk or hashtag) and corresponding p-values.

## Results

### ODE exposure induces differential gene expression in mouse brain microglia, leading to upregulation in pathways associated with biological processes (BP), molecular function (MF)

A comprehensive transcriptomic analysis was performed on microglia isolated from mice to identify differentially expressed genes (DEGs). The analysis revealed 19 DEGs in the ODE-exposed group, each showing a minimum 2-fold change and a *p*-value ≤ 0.05 (Fig. 1). Among these, 17 genes were upregulated, and 2 genes were downregulated in response to ODE exposure (Fig. 1A–B). Refer to Supplementary Table 1 for details on DEGs. Pathway enrichment analysis of these DEGs was conducted using DAVID analysis (The Database for Annotation, Visualization, and Integrated Discovery). The significantly enriched pathways from biological processes (BP), molecular functions (MF), and cellular components (CC) were clustered and visualized (Fig. 1C). Notably, the enriched pathways primarily belonged to BP and MF categories in response to ODE exposure. Further details on these enriched pathways are provided in Supplementary Table 2.

**Figure 1.**
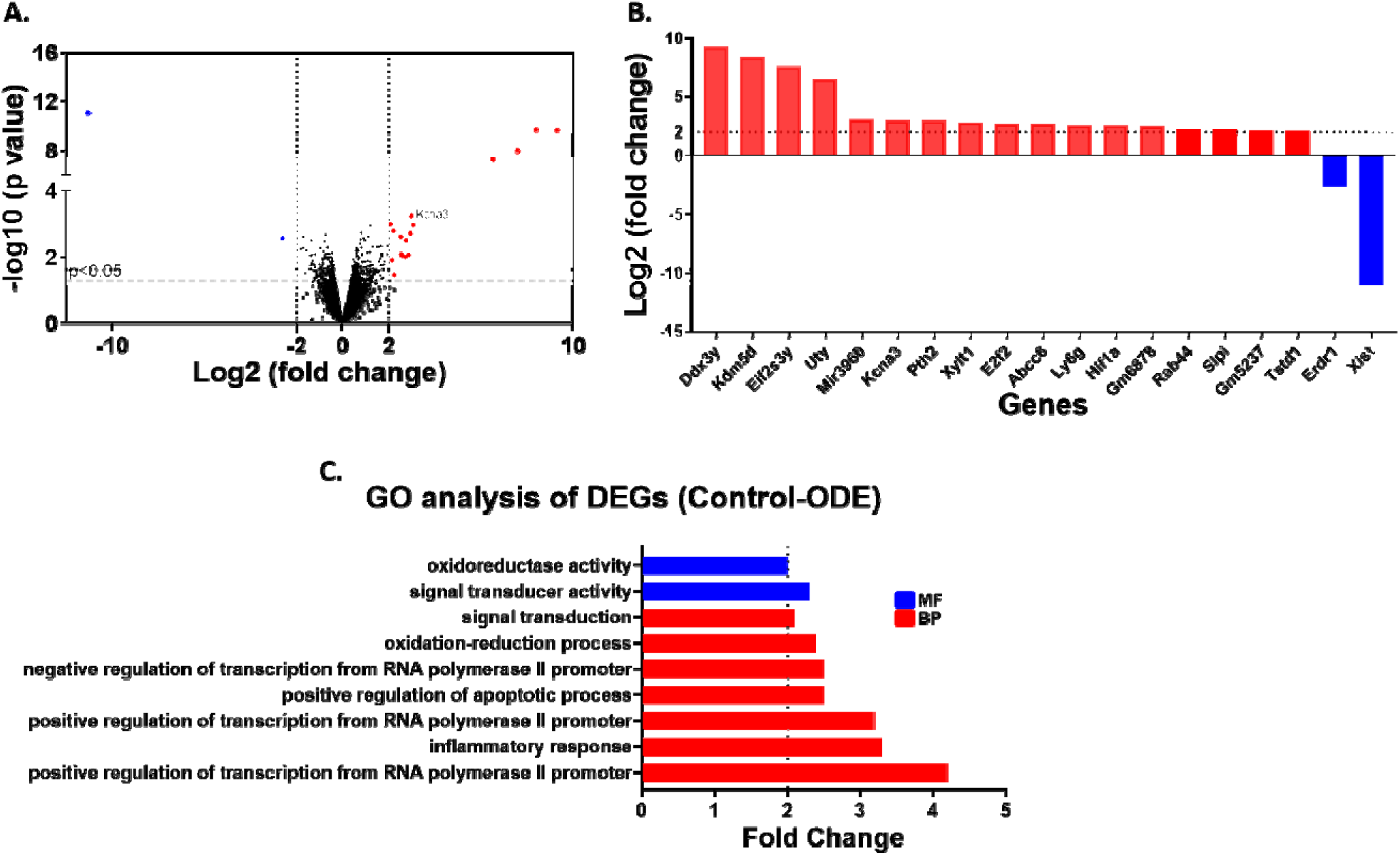
Differential gene expression (DEGs) analysis and enriched pathways based onGene ontology (GO) in isolated microglia following RNA-seq. After five weeks of treatment, brains were collected from mice, and the microglial fraction was isolated. Total RNA was extracted from the isolated microglia and processed for RNA-seq analysis. Differentially expressed genes (DEGs) were identified based on statistical significance (*p* ≤ 0.05) and a log_₂_fold-change > 2. A volcano plot and bar graph illustrate these DEGs in ODE-exposed mouse brains (A-B). Pathway enrichment analysis was performed using DAVID, focusing on gene ontology (GO) terms. Significantly enriched pathways associated with biological processes (BP) and molecular functions (MF), meeting the criteria of *p* ≤ 0.05 and log_₂_ fold-change > 2, are plotted in (C). The analysis included *n*=3 male mice.

### DEGs were validated for GO analysis through qPCR following RNA-seq of isolated microglia

A total of 40 genes were selected for validation using qPCR. Of these, some were identified as differentially expressed genes (DEGs) from the RNA-seq analysis, while others were chosen based on pathway enrichment analysis, where specific pathways indicated significant expression of additional genes (Fig. 2). Among these, six genes (*Hif1*α, *Hif2*α, *E2f2*, *Ddx3y*, *Kcna3*, and *Xylt1*) consistently showed significant validation results, with a log_₂_fold-change > 2 compared to the control group. Refer to Supplementary Table 3 for the list of 40 genes.

**Figure 2.**
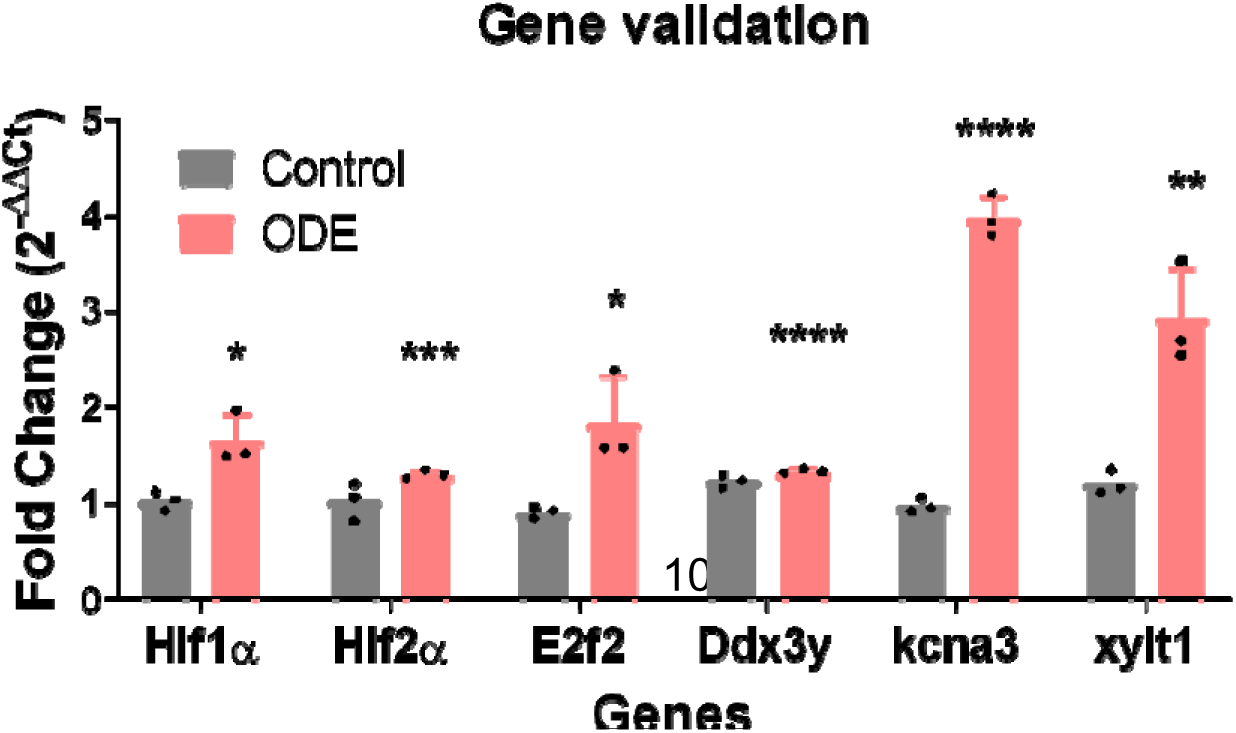
Validation of DEGs identified through RNA-seq and DAVID analysis using qPCR. After five weeks of exposure, RNA was extracted from isolated microglia, treated with DNase, and processed to remove rRNA for gene validation through qPCR analysis. Out of the 40 genes selected for validation (Supplementary Table 3), six genes demonstrated a significant log_₂_ fold-change > 2 compared to the control group. The validated genes include Hypoxia-inducible factor 1α (*Hif1*α), Hypoxia-inducible factor 2α (*Hif2*α), E2F transcription factor 2 (*E2f2*), DEAD (Asp-Glu-Ala-Asp) box polypeptide 3, Y-linked (*Ddx3y*), Potassium voltage-gated channel (*Kcna3*), and Xylosyltransferase 1 (*Xylt1*). The analysis was performed using *n*=3 male mice. Statistical significance is indicated by (* for ODE exposure effect), with detailed *p*-values and statistical tests provided in Table 7.

**Table 7.** Details of statistical analysis applied for each experiment/figure.

| Figur | Panel | Test |
| --- | --- | --- |
| 1 | A-B | one-way ANOVA; Tukey's post-hoc, n=3 mice |
| 2 |  | one-way ANOVA; Tukey's post-hoc, n=3 mice |
| 3 | A-C | one-way ANOVA; Tukey's post-hoc, n=3 mice |
| 4 | A&D | two-way ANOVA; Tukey's post-hoc; n=3 wells (4-5 slices/ well) (6 well plate) |
|  | B&E | two-way ANOVA; Tukey's post-hoc; n=3 wells (24 well plate) |
|  | C&F | two-way ANOVA; Tukey's post-hoc; n=3 wells (24 well plate) |
| 5 | B | two-way ANOVA; Tukey's post-hoc; n=3 wells (4-5 slices/ well) (6 well plate) |
|  | D&F | two-way ANOVA; Tukey's post-hoc; n=3 wells (24 well plate) |
| 6 | B | two-way ANOVA; Tukey's post-hoc; n=3 wells (4-5 slices/ well) (6 well plate) |
|  | D | two-way ANOVA; Tukey's post-hoc; n=3 wells (24 well plate) |
| 7 | B | two-way ANOVA; Tukey's post-hoc; n=3 wells (4-5 slices/ well) (6 well plate) |
|  | D&F | two-way ANOVA; Tukey's post-hoc; n=3 wells (24 well plate) |
| 8 | A | two-way ANOVA; Tukey's post-hoc; n=3 wells (4-5 slices/ well) (6 well plate) |
|  | B-C | two-way ANOVA; Tukey's post-hoc; n=3 wells (24 well plate) |
| 9 | A-B | two-way ANOVA; Tukey's post-hoc; n=3 wells (4-5 slices/ well) (6 well plate) |
| 10 | A-B | two-way ANOVA; Tukey's post-hoc; n=3 wells (24 well plate) |
| 11 | H,I,J | two-way ANOVA; Tukey's post-hoc; n=3 wells (24 well plate) |

### ODE exposure increased the expression of Kv1.3, NOX2, and phosphorylated-p38 MAPK (p-p38 MAPK) in mouse brain following ODE exposure

After five weeks of exposure, total protein was isolated from mouse brains and analyzed for Kv1.3, NOX2, and p-p38 MAPK protein expression using western blot analysis (Fig. 3). Compared to controls, five weeks of ODE exposure resulted in increased expression levels of Kv1.3, NOX2, and p-p38 MAPK in mouse brains (Fig. 3A–C).

**Figure 3.**
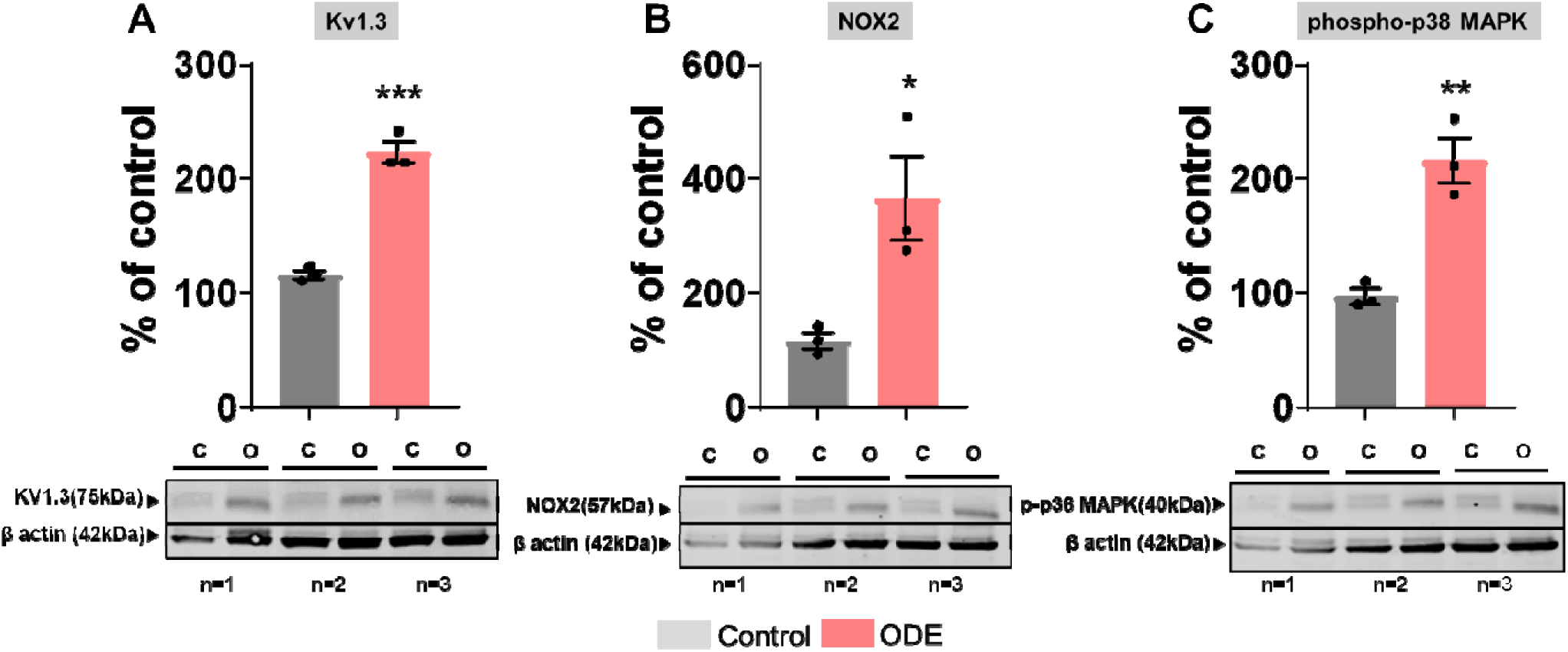
Kv1.3, NOX2, and p-p38 MAPK14 protein expression in mice brains. Western blot analysis was performed on whole-cell lysates from freshly dissected mouse brains to evaluate the expression levels of Kv1.3 (A), NOX2 (B), p-p38 MAPK14 (C), and β-actin (housekeeping protein). The detected bands corresponded to molecular weights of 75 kDa (Kv1.3), 57 kDa (NOX2), 40 kDa (p-p38 MAPK14), and 42 kDa (β-actin). Densitometric analysis of normalized protein bands revealed that ODE exposure significantly increased NOX2 protein levels in mouse brains compared to controls (*n=3*/group, male mice). Statistical significance is indicated by (* for ODE exposure effect). Detailed *p-*values and statistical tests are provided in Table 7.

### ODE induces *kcna3* and *MAPK14* expression in BSCs, BV2 microglia and primary microglia

After five weeks of treatment, total RNA (free from genomic DNA) was extracted from BSCs, BV2 microglia, and primary microglia and subsequently analyzed using qPCR (Fig. 4). The results revealed a significant increase in the gene expression of *kcna3* (Fig. 4A-C) and *MAPK14* (Fig. 4D-F) in BSCs, BV2 microglia, and primary microglia compared to the control group. Notably, treatment with PAP-1 significantly reduced the expression levels of *kcna3* (Fig. 4A-C) and *MAPK14* (Fig. 4D-F) in ODE-treated BSCs, BV2 microglia, and primary microglia.

**Figure 4.**
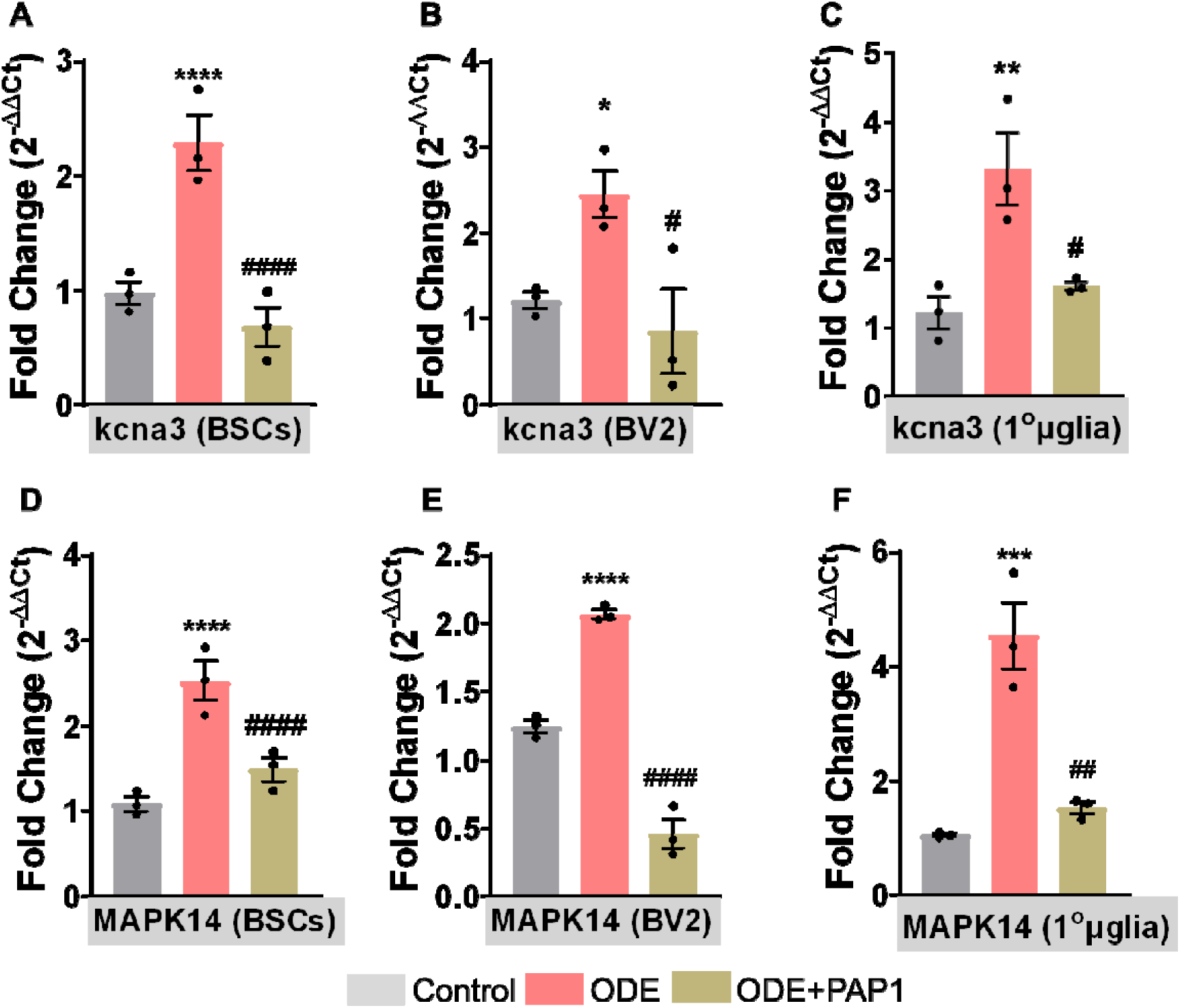
Regulation of *kcna3* and *MAPK14* gene expression by PAP-1 in ODE-treated BSCs, BV2 microglia, and primary microglia. qPCR analysis was performed on total RNA (free from genomic DNA) extracted from BSCs, BV2 microglia, and primary microglia (referred to as 1°µglia) to evaluate the expression of *Kcna3* (A) and *MAPK14* (B) genes. Specifically designed primers used for this analysis are listed in Table 4. The ODE-exposed group exhibited significantly elevated mRNA expression of *Kcna3* (A-C) and *MAPK14* (D-F) compared to the control group. In contrast, the PAP-1 co-treatment group showed significantly reduced mRNA expression levels of both *Kcna3* (A-C) nd *MAPK14* (D-F) in BSCs, BV2 microglia, and primary microglia. The sample size included *n*=3 mouse pups for BSC and primary microglia; and *n*=3 independent cultures for BV2 microglia (biological replicates). Statistical significance is indicated by (* for ODE exposure effect and # for PAP-1 treatment effect), with corresponding *p*-values and statistical tests detailed in Table 7.

### ODE induces Kv1.3 expression in BSCs, BV2 microglia and primary microglia

Following treatments with ODE and PAP-1, BSCs, BV2 microglia, and primary microglia were analyzed for cellular expression of Kv1.3 protein using immunostaining (Fig. 5). Quantitative analysis of fluorescent images in BSCs (Fig. 5Ai–Aiii, B), BV2 microglia (Fig. 5Ci–Ciii, D), and primary microglia (Fig. 5Ei–Eiii, F) revealed mean Kv1.3 expression per cell. ODE exposure resulted in a significant increase in Kv1.3 protein expression in BSCs (Fig. 5B), BV2 microglia (Fig. 5D), and primary microglia (Fig. 5F) compared to the control group. However, co-treatment with PAP-1 significantly reduced ODE-induced Kv1.3 protein expression in BSCs (Fig. 5A–B), BV2 microglia (Fig. 5C–D), and primary microglia (Fig. 5E–F).

**Figure 5.**
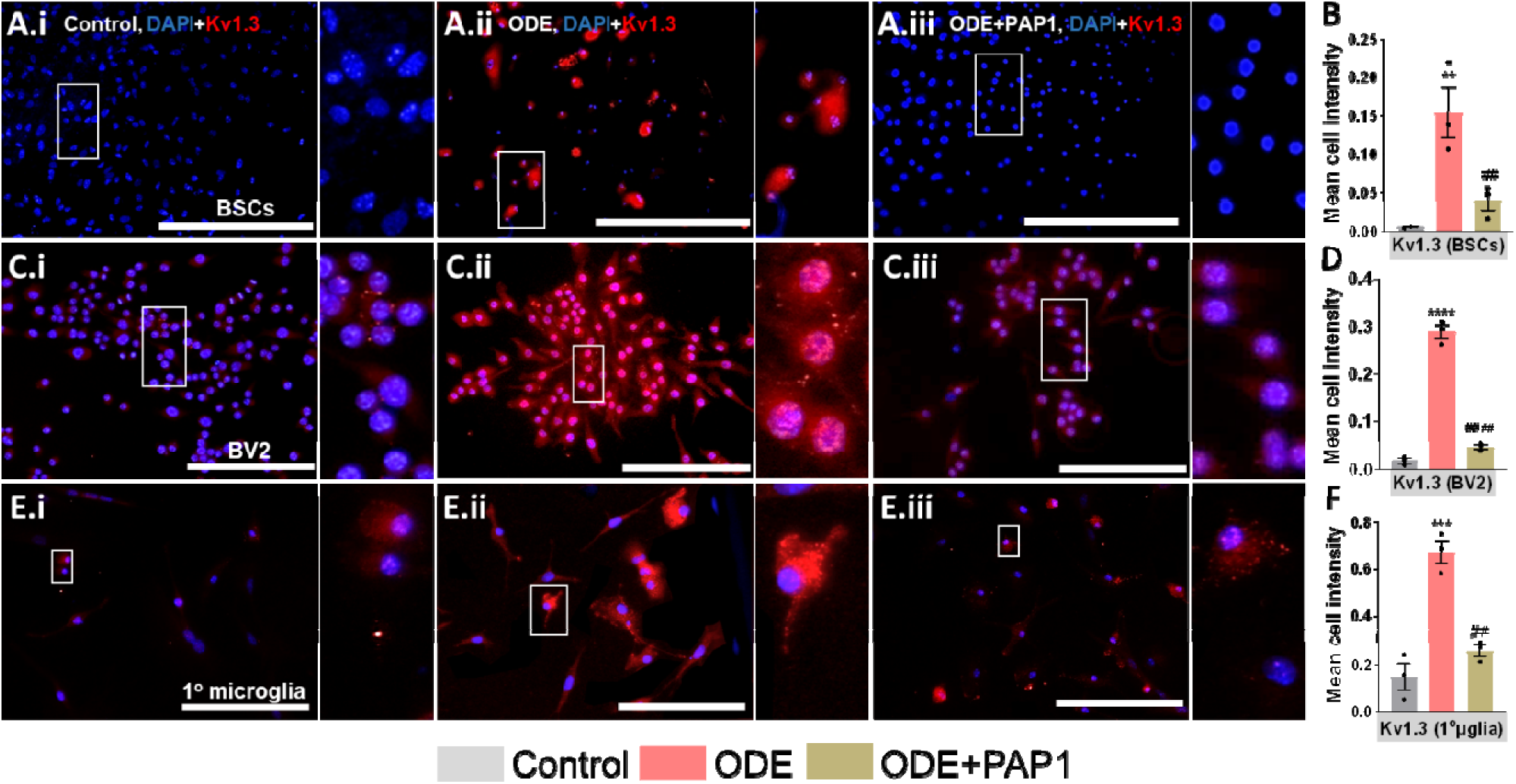
IHC and ICC analysis of Kv1.3 expression in ODE-treated BSCs, BV2 microglia, and primary microglia with PAP-1 treatment effects. Following the treatments (Table 1), BSCs (Ai-iii), BV2 microglia (Bi-iii), and primary microglia (referred to as 1°µglia) (Ci-iii) were immunostained for Kv1.3 (Cy3, red). The antibodies used for immunostaining are detailed in Table 5. Nuclei were counterstained with DAPI (blue) to visualize and count the number of cells in each field. To specifically identify microglia expressing Kv1.3 in BSCs, co-staining with Iba1 (a microglia marker) was performed. Images of Kv1.3+Iba1 co-staining are provided in Supplementary Figure 1. Quantification of immunohistochemistry (IHC) and immunocytochemistry (ICC) staining for Kv1.3 expression was carried out as described in the *Methods* section. Compared to the control group, ODE-exposed groups showed a significant increase in Kv1.3 staining intensity in BSCs (B), BV2 microglia (D), and primary microglia (referred to as 1°µglia) (F). Notably, PAP-1 treatment significantly reduced Kv1.3 expression in BSCs (A–B), BV2 microglia (C–D), and primary microglia (E–F), as shown in the representative images and quantitative analysis. The sample size included *n*=3 mouse pups for BSC and primary microglia; and *n*=3 independent cultures for BV2 microglia (biological replicates). Statistical significance is indicated by an * for the ODE exposure effect and # for the PAP-1 treatment effect. The scale bar represents 50 µm. Details of *p*-values and statistical tests are provided in Table 7.

### ODE induces NOX2 expression in BSCs and primary microglia

Following treatments with ODE and PAP-1, BSCs and primary microglia were analyzed for cellular expression of NOX2 protein using immunostaining (Fig. 6). Quantitative analysis of fluorescent images revealed mean NOX2 expression per cell in BSCs (Fig. 6Ai–Aiii, B) and primary microglia (Fig. 6Ci–Ciii, D). ODE exposure resulted in a significant increase in NOX2 protein expression in BSCs (Fig. 6B) and primary microglia (Fig. 6D) compared to the control group. However, co-treatment with PAP-1 significantly reduced ODE-induced NOX2 protein expression in BSCs (Fig. 6A–B) and primary microglia (Fig. 6C–D).

**Figure 6.**
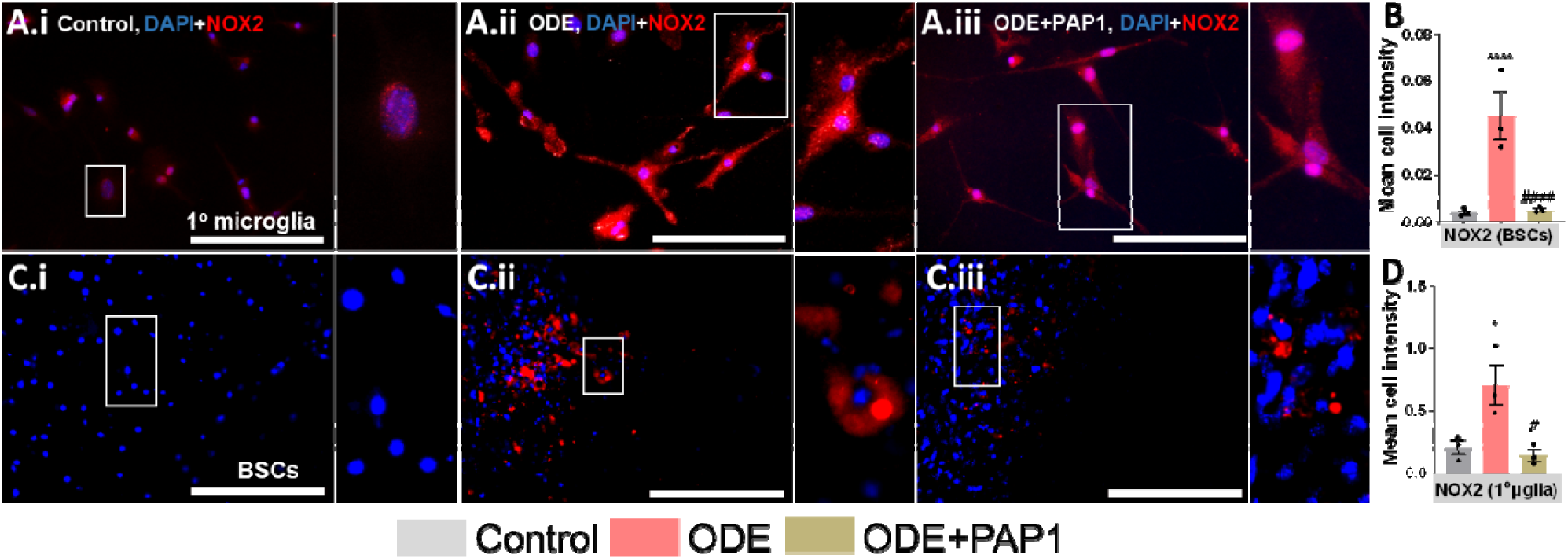
Immunohistochemical analysis of NOX2 expression in ODE-treated BSCs, and primary microglia with PAP-1 treatment effects. Following the treatments (Table 1), BSCs (Ai-iii), and primary microglia (referred to as 1°µglia) (Ci-iii) were immunostained for NOX2 (Cy3, red). The antibodies used for immunostaining are detailed in Table 5. Nuclei were counterstained with DAPI (blue) to visualize and count the number of cells in each field. To specifically identify microglia expressing Kv1.3 in BSCs, co-staining with Iba1 (a microglia marker) was performed. Images of Kv1.3+Iba1 co-staining are provided in Supplementary Figure 1. Quantification of immunohistochemistry (IHC) and immunocytochemistry (ICC) staining for NOX2 expression was carried out as described in the *Methods* section. Compared to the control group, ODE-exposed groups showed a significant increase in NOX2 staining intensity in BSCs (B), and primary microglia (referred to as 1°µglia) (D). Notably, PAP-1 treatment significantly reduced NOX2 expression in BSCs (A–B), and primary microglia (C–D), as shown in the images and quantitative analysis. The sample size included *n*=3 mouse pups for BSC and primary microglia. Statistical significance is indicated by (* for ODE exposure effect and # for PAP-1 treatment effect). The scale bar represents 50 µm. Details of *p*-values and statistical tests are provided in Table 7.

### ODE induces p-p38 MAPK expression in BSCs, BV2 microglia and primary microglia

Following treatments with ODE and PAP-1, BSCs, BV2 microglia, and primary microglia were analyzed for cellular expression of p-p38 MAPK protein using immunostaining (Fig. 7). Quantitative analysis of fluorescent images in BSCs (Fig. 7Ai–Aiii, B), BV2 microglia (Fig. 7Ci–Ciii, D), and primary microglia (Fig. 7Ei–Eiii, F) revealed mean p-p38 MAPK expression per cell. ODE exposure resulted in a significant increase in p-p38 MAPK protein expression in BSCs (Fig. 7B), BV2 microglia (Fig. 7D), and primary microglia (Fig. 7F) compared to the control group. However, co-treatment with PAP-1 significantly reduced ODE-induced p-p38 MAPK protein expression in BSCs (Fig. 7A–B), BV2 microglia (Fig. 7C–D), and primary microglia (Fig. 7E–F).

**Figure 7.**
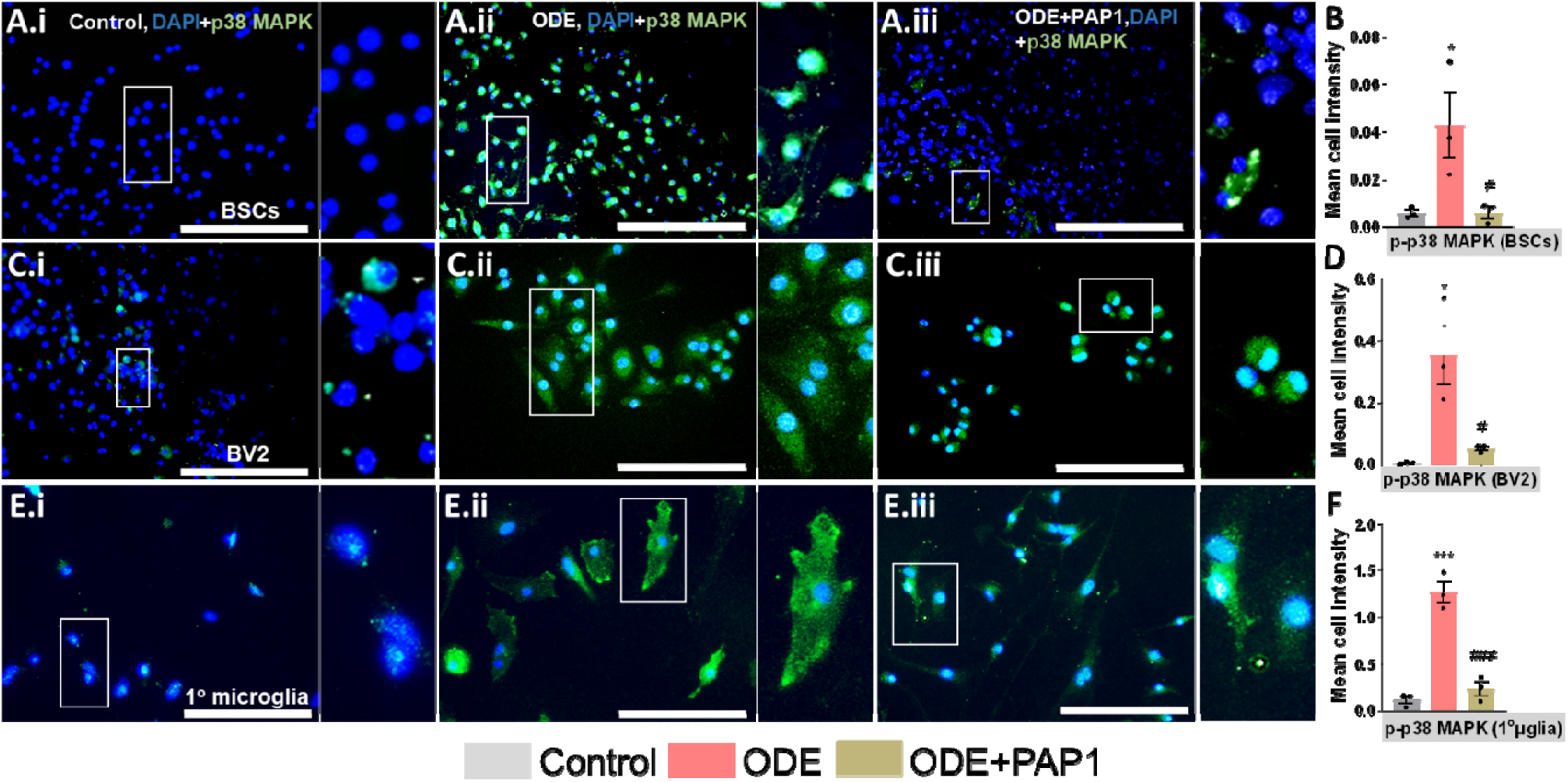
Immunohistochemical analysis of p-p38 MAPK expression in ODE-treated BSCs, BV2 microglia, and primary microglia with PAP-1 treatment effects. Following the treatments (Table 1BSCs (Ai-iii), BV2 microglia (Bi-iii), and primary microglia (referred to as 1°µglia) (Ci-iii) were immunostained for p-p38 MAPK (FITC, green). The antibodies used for immunostaining are detailed in Table 5. Nuclei were counterstained with DAPI (blue) to visualize and count the number of cells in each field. To specifically identify microglia expressing p-p38 MAPK in BSCs, co-staining with Iba1 (a microglia marker) was performed. Images of p-p38 MAPK +Iba1 co-staining are provided in Supplementary Figure 1. Quantification of immunohistochemistry (IHC) and immunocytochemistry (ICC) staining for p-p38 MAPK expression was carried out as described in the Methods section. Compared to the control group, ODE-exposed groups showed a significant increase in p-p38 MAPK staining intensity in BSCs (B), BV2 microglia (D), and primary microglia (referred to as 1°µglia) F). Notably, PAP-1 treatment significantly reduced p-p38 MAPK expression in BSCs (A–B), BV2 microglia (C–D), and primary microglia (E–F), as shown in the representative images and quantitative analysis. The sample size included *n*=3 mouse pups for BSC and primary microglia; and *n*=3 independent cultures for BV2 microglia (biological replicates). Statistical significance is indicated by (* for ODE exposure effect and # for PAP-1 treatment effect). The scale bar represents 50 µm. Details of *p*-values and statistical tests are provided in Table 7.

### ODE exposure induces nitrite secretion in BSCs, BV2 microglia, and primary microglia

Cell culture supernatants from BSCs, BV2 microglia, and primary microglia were analyzed for nitrite levels (Fig. 8). ODE exposure resulted in a significant increase in secreted nitrite levels in the supernatants of BSCs (Fig. 8A), BV2 microglia (Fig. 8B), and primary microglia (Fig. 8C) compared to controls. However, co-treatment with PAP-1 significantly reduced nitrite levels in the culture supernatants of ODE-exposed BSCs, BV2 microglia, and primary microglia.

**Figure 8.**
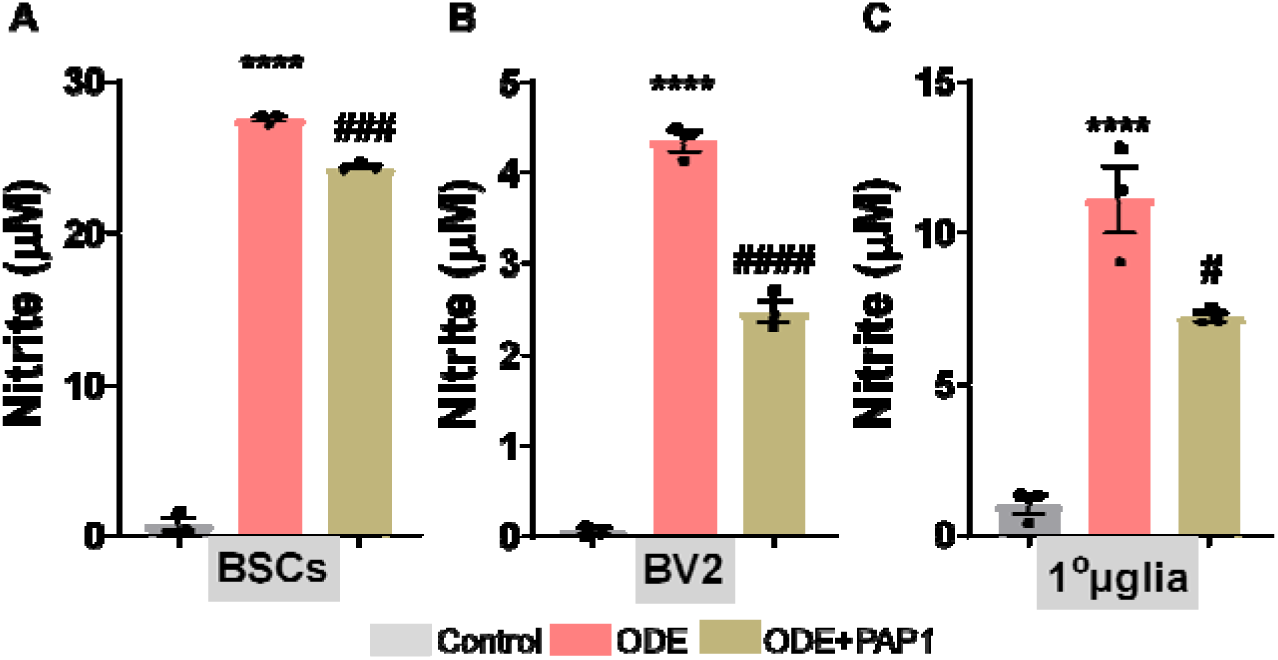
Griess assay results for culture supernatant of BSCs, BV2 microglia, and primary microglia. The Griess Assay Kit was used to measure the concentration of secreted nitrite in the\ culture supernatants. ODE-exposed groups showed a significant increase in nitrite levels in the supernatants of BSCs (A), BV2 microglia (B), and primary microglia (referred to as 1°µglia) (C) compared to control groups. Co-treatment with PAP-1 significantly reduced nitrite levels in the supernatants of BSCs, BV2 microglia, and primary microglia. The results are presented as µM of nitrite in the supernatant. The sample size included *n*=3 mouse pups for BSC and primary microglia; and *n*=3 independent cultures for BV2 microglia (biological replicates). Statistical significance is indicated by (* for ODE exposure effect and # for PAP-1 treatment effect). Further details on *p*-values and statistical tests are provided in Table 7.

### ODE exposure induces pro-inflammatory cytokine secretion in BSCs

Culture supernatants from BSCs were collected and analyzed for pro-inflammatory cytokine levels using ELISA (Fig. 9). ODE exposure resulted in a significant increase in the secretion of TNF-α (Fig. 9A) and IL-6 (Fig. 9B) into the culture supernatants of BSCs compared to the control group. Co-treatment with PAP-1 significantly reduced the levels of IL-6 but not TNF-α in the culture supernatants.

**Figure 9.**
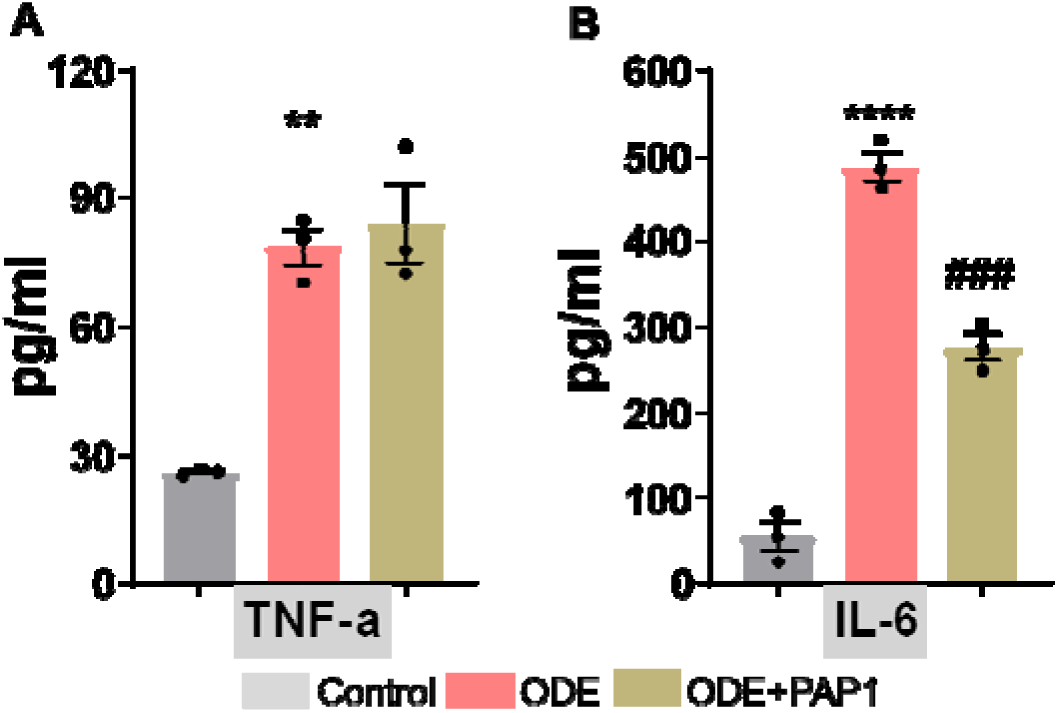
Cytokine analysis of BSCs culture supernatants. Commercially available cytokine kits were used to measure pro-inflammatory cytokine levels in the culture supernatants of BSCs. ODE-exposed groups showed a significant increase in secreted TNF-α (A) and IL-6 (B) levels compared to control groups. PAP-1 co-treatment significantly reduced the secretion of IL-6 (B) in BSCs. The results are expressed in pg/mL. The sample size included *n=*3 mouse pups. Statistical significance is indicated by (* for ODE exposure effect and # for PAP-1 treatment effect). Detailed *p*-values and statistical tests are provided in Table 7.

### ODE exposure induced cytokine secretion in BV2 microglia and primary microglia

Cell culture supernatants from BV2 microglia and primary microglia were collected and analyzed using multiplex cytokine assays (Fig. 10). In BV2 microglia, ODE exposure resulted in a significant increase in the levels of TNF-α, IL-6, IL-1β, and IL-17A (Fig. 10A). In primary microglia, elevated levels of TNF-α, IL-6, IL-1β, IL-10, IL-17A, and IFNγ were observed following ODE exposure (Fig. 10B). Notably, PAP-1 treatment significantly reduced ODE-induced TNF-α secretion in BV2 microglia supernatants only (Fig. 10A).

**Figure 10.**
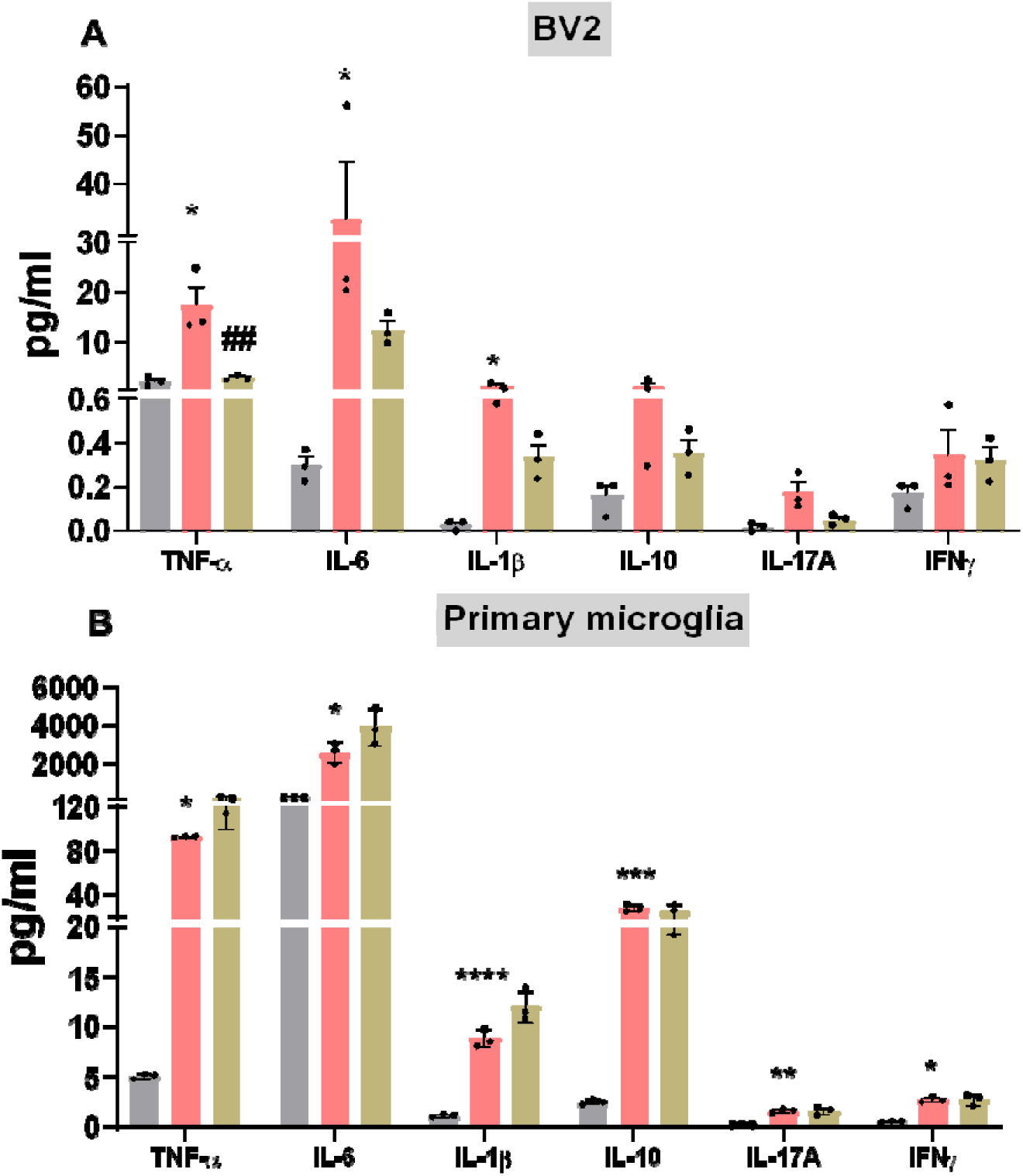
Cytokine profile in ODE-exposed BV2 and primary microglia with PAP-1 treatment effects. A customized MILLIPLEX® Mouse Cytokine/Chemokine Kit was used to analyze cytokine levels in culture supernatants from BV2 microglia and primary microglia. Compared to control groups, ODE exposure significantly increased the levels of TNF-α, IL-6, IL-1β, and IL-17A in BV2 microglia (A) and TNF-α, IL-6, IL-1β, IL-10, IL-17A, and IFNγ in primary microglia (B). PAP-1 co-treatment significantly reduced TNF-α levels in BV2 microglia supernatants (A) but did not significantly affect other cytokines in either cell type. The results are expressed in pg/mL. The sample size included *n*=3 mouse pups for primary microglia and *n*=3 independent cultures for BV2 microglia (biological replicates). Statistical significance is indicated by (* for ODE exposure effect and # for PAP-1 treatment effect). Details of *p*-values and statistical tests are provided in Table 7.

### ODE exposure increases electrical capacitance without affecting Kir2.1 and Kv1.3 currents in primary microglia

Primary microglia were treated with either unstimulated medium (DMEM with 2% FBS), LPS (100 ng/mL) as a positive control, or ODE (1% v/v) for 48 hours. Whole-cell patch-clamp recordings were performed on individual microglia at 48 hours following treatment. Light microscopic evaluation revealed a notable increase in microglial cell size with an ameboid morphology after LPS and ODE stimulation compared to unstimulated controls (Fig. 11A-C), indicating an activated state. Interestingly, while LPS induced Kv1.3 currents and decreased Kir2.1 currents as expected, organic dust did not induce significant Kv1.3 currents on the plasma membrane or decrease Kir2.1 currents (Fig.11D-J). ODE certainly activated microglia based on the cell shape and the increase in electrical capacitance.

**Figure 11.**
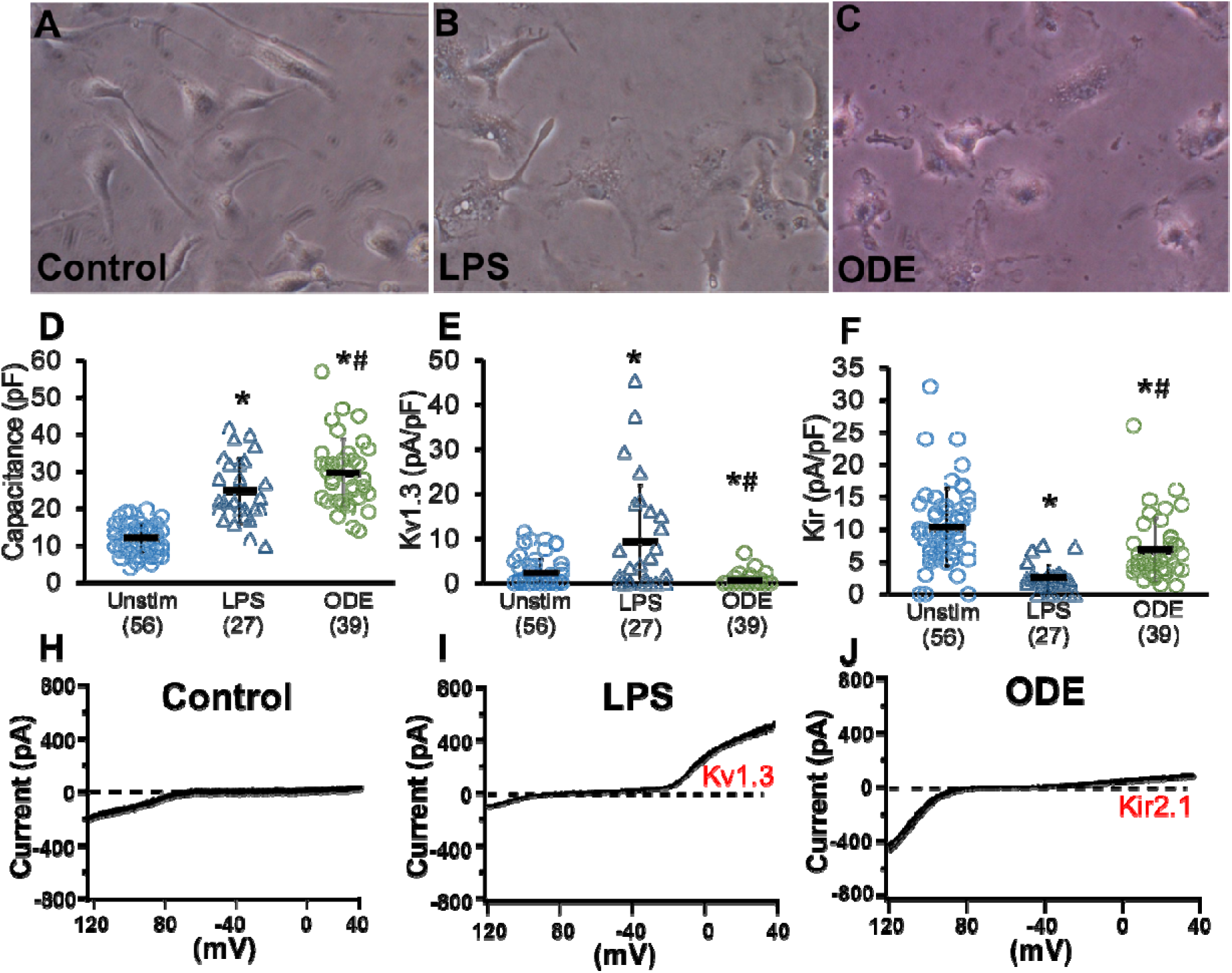
Electrophysiological and morphological changes in primary microglia following LPS and ODE stimulation. Primary microglia were either treated with vehicle (DMEM in 2% FBS), LPS (100 ng/mL), or ODE (1% v/v), and whole cell patch clamp recordings were taken at 48 hours. Light microscopic microphotographs are shown to visualize structural changes in microglia (A-C). The rise in Kv1.3 and Kir2.1 currents following DMEM with 2%FBS (control). LPS (100ng/mL) and ODE (1% w/v) after 48-hour treatment are shown (D-F). *n=*3 mouse pups; * ODE exposure effect (LPS or ODE). *p-*values and statistical tests are listed in Table 7.

## Discussion

Agricultural regions in the United States have been associated with exposure to agricultural contaminants that may play a role in initiating and perpetuating neuroinflammatory cycles, ultimately contributing to neurodegeneration (Wright Willis et al., 2010) (Arora et al., 2021). Building on our previous published research (Massey et al., 2019; Massey et al., 2021a; Massey et al., 2022), this study aimed to investigate the mechanisms by which inhaled organic dust extract (ODE) induces sustained neurotoxic signaling. Chronic neuroinflammation often accompanies neurodegenerative diseases and involves a dynamic interplay among various cell types, including neurons, microglia, astrocytes, and peripheral immune cells, and is mediated through a wide range of signaling molecules such as cytokines, chemokines, reactive oxygen species (ROS), and damage-associated molecular patterns (DAMPs). This complex interplay not only makes it difficult to understand the disease process but also poses a challenge to developing effective therapeutic strategies. Given the complexity and heterogeneity of the underlying pathways, a comprehensive transcriptomic analysis at the genetic level can provide significant insights. Transcriptomic analysis can identify changes in the global gene expression across various cell types in the brain and pinpoint the central regulatory proteins serving as potential therapeutic targets.

The findings from our current study provide new insights into the transcriptomic and molecular pathways underlying ODE-induced neuroinflammation and neurodegeneration. RNA sequence analysis revealed that ODE exposure significantly upregulates the expression of the *kcna3* gene, which encodes the Kv1.3 potassium channel. The voltage-gated potassium channel Kv1.3 is pivotal in microglial activation and the subsequent neuroinflammatory response (Rangaraju et al., 2017). Upregulation of Kv1.3 in microglia has been observed in various neurodegenerative conditions, including Alzheimer’s and Parkinson’s diseases (Fordyce et al., 2005; I. Maezawa et al., 2018; Sarkar et al., 2020). This upregulation promotes the production of pro-inflammatory cytokines and reactive oxygen species (ROS), exacerbating neuronal damage. Furthermore, inhibition of Kv1.3 has been shown to attenuate microglial activation and reduce neuroinflammation, highlighting its potential as a therapeutic target (Sarkar et al., 2020). We chose the kcna3 gene owing to its crucial role in microglial function and its importance in neuroinflammation and neurodegeneration. It encodes the Kv1.3 potassium channel, regulating microglial activation, proliferation, and cytokine production. Kv1.3 dysregulation is linked to chronic neuroinflammation, mitochondrial dysfunction, and oxidative stress, making it a key therapeutic target. Investigating the role of *kcna3* may help link gene expression changes to cellular and molecular dysfunctions, providing insights into microglial pathology and neurotoxicity.

In this study, the upregulation of *kcna3* was accompanied by increased levels of the *MAPK14* gene. MAPK14 encodes p38α MAPK, a key member and prototypical representative of the p38 MAPK family. This kinase plays a central role in cellular responses to stress and inflammation, acting as a critical regulator of signaling pathways involved in immune responses, apoptosis, and cytokine production (B. Canovas & A. R. Nebreda, 2021). Activation of p38 MAPK in microglia has been linked to producing pro-inflammatory mediators, contributing to neurotoxicity. Selective inhibition of p38α MAPK has been demonstrated to alleviate neuroinflammation and associated neuropathology, underscoring its significance in neurodegenerative diseases (Gee et al., 2020). The interaction between Kv1.3 channels and the p38 MAPK pathway suggests a complex signaling network that modulates microglial activation and neuroinflammation. Studies indicate that Kv1.3 may influence p38 MAPK activity and vice versa, creating a feedback loop that amplifies inflammatory responses. Disruption of this interaction through targeted inhibition could provide a multifaceted approach to mitigating neuroinflammatory processes (Rangaraju et al., 2017). This signaling cascade is critical for inflammatory responses and has been shown to modulate Kv1.3 levels via Fyn Kinase-mediated p38 MAPK phosphorylation and NF-κB pathway activation (Panicker et al., 2015; Sarkar et al., 2020). Additionally, p-p38 MAPK enhances microglial neurotoxicity by promoting Kv1.3 expression and the generation of peroxynitrite, a neurotoxic compound formed from the interaction of reactive oxygen species (ROS) and nitrite (Jianuo Liu et al., 2012), a neurotoxic compound produced from the reaction of ROS and nitrite (Fordyce et al., 2005) (Torreilles et al., 1999) In our study, brain tissues from mice exposed to ODE for five weeks demonstrated significant increases in Kv1.3, p-p38 MAPK, and NOX2, implicating oxidative stress as a key driver of neuroinflammatory signaling.

To assess the translational relevance of our findings, we utilized three experimental models: brain slice cultures (BSCs), a microglial cell line (BV2 microglia), and primary microglial cultures. Following exposure to organic dust extract (ODE), there was a significant elevation in the expression of *kcna3* and *MAPK14* genes. Immunofluorescence imaging further demonstrated increased levels of microglial Kv1.3 and phosphorylated p38 MAPK across all models.

Pharmacological inhibition of Kv1.3 has shown promise in reducing microglial-mediated neuroinflammation. Selective blockers, such as PAP-1 and ShK-223, have been effective in attenuating microglial activation and decreasing the production of pro-inflammatory cytokines. These findings support the potential of Kv1.3 inhibitors as therapeutic agents in other neuroinflammatory and neurodegenerative disorders (Rangaraju et al., 2017). The critical role of Kv1.3 in the ODE exposure model was also confirmed using PAP-1 in our study. Co-treatment of PAP-1 with ODE significantly reduced the expression of *kcna3* and *MAPK14* genes, accompanied by a marked decrease in Kv1.3 and p-p38 MAPK protein levels. These findings highlight the central role of Kv1.3 upregulation in ODE-induced neuroinflammation and demonstrate that PAP-1-mediated inhibition effectively mitigates these neuroinflammation in our models. Similarly, targeting the p38 MAPK pathway could also offer a potential therapeutic target in combination with Kv1.3. Selective p38α MAPK inhibitors have been shown to alleviate neuropathology and improve cognitive function in models of neurodegenerative diseases. The combined inhibition of Kv1.3 channels and the p38 MAPK pathway represents a promising strategy for controlling microglial activation and reducing neuroinflammation (Garrett & Kyriacou, 1988).

Overwhelming oxidative stress, caused by an imbalance between ROS and antioxidants, is a key factor in driving neuroinflammation resulting in neurodegenerative diseases. Microglia, contribute to this process by producing ROS through enzymes such as NOX2. Excessive ROS leads to cellular damage and exacerbates inflammation, further harming neurons (Houldsworth, 2024). The MAPK pathway plays a central role in this process, as oxidative stress activates it, resulting in the production of pro-inflammatory cytokines. This creates a feedback loop where oxidative stress and inflammation perpetuate each other, implicated in diseases such as Alzheimer’s and Parkinson’s. Elevated levels of p38 MAPK and oxidative stress markers are observed in these two conditions (Bernhardi, 2009).

The relationship between Kv1.3 channel activity and oxidative stress was further validated in our ODE exposure model by examining NOX2, a key enzyme responsible for ROS production. NOX2 is a central player in the oxidative stress pathways linked to microglial activation and subsequent neuronal damage. Consistent with our previous findings (Massey et al., 2019), ODE exposure significantly increased NOX2 protein expression and nitrite release in both primary microglia and brain slice cultures (BSCs). These increases were accompanied by elevated levels of Kv1.3 and p-p38 MAPK, suggesting a mechanistic link in which Kv1.3 channel activity drives oxidative stress. The resulting oxidative environment facilitates the formation of peroxynitrite, a neurotoxic compound contributing to neuronal damage. This interaction underscores the intricate interplay between Kv1.3 function and the oxidative stress pathways that exacerbate inflammation and neuronal injury. The effectiveness of PAP-1 in reducing these effects reinforces the role of Kv1.3 not only as a biomarker for microglial activation but also as a potential therapeutic target for mitigating neuroinflammatory damage in the ODE exposure model. By targeting Kv1.3, it may be possible to interrupt the vicious cycle of ROS production, microglial over activation, and neuronal degeneration, offering a novel therapeutic intervention in neuroinflammatory diseases. Further studies exploring the precise molecular mechanisms linking Kv1.3 activity with oxidative stress pathways will likely provide deeper insights into its role in neuroinflammation and help refine therapeutic strategies (Jembrek, 2024).

We investigated inflammatory cytokines as markers of microglial activation to understand the inflammatory landscape induced by ODE exposure. Our findings showed that ODE exposure significantly increased the release of pro-inflammatory cytokines such as TNF-α, IL-6, IL-1β, and IL-17A as previously shown (Massey et al., 2019; Massey et al., 2021a; Massey et al., 2022). In primary microglial cultures, a broader spectrum of cytokines, including IL-10 and IFNγ, were observed, indicating a robust inflammatory response and heightened microglial activation. PAP-1, a Kv1.3 inhibitor, reduced IL-6 but not the TNF-α levels, highlighting Kv1.3’s role in cytokine regulation but suggesting that other pathways may also be involved. The presence of IL-10 suggests simultaneous activation of pro- and anti-inflammatory pathways. The complexity of ODE-induced inflammation underscores the need to explore other pathways, such as TLRs, MAPKs, or inflammasomes, to better understand and mitigate neuroinflammation. Future studies should focus on identifying additional therapeutic targets to complement Kv1.3 inhibition in ODE exposure models.

Interestingly, whole-cell patch-clamp experiments revealed no enhancement of Kv1.3 currents on the cell surface despite increased intracellular Kv1.3 protein levels. This suggests that Kv1.3 primarily accumulates intracellularly rather than being inserted into the plasma membrane, indicating a disruption in protein trafficking and membrane localization. Such disruptions could stem from damage to the endoplasmic reticulum (ER) and mitochondria, as previously observed in our studies where ODE exposure damaged ER and mitochondria and negatively affected mitochondrial bioenergetics (Massey et al., 2021a) (Bowen et al., 2024).

Incorporating findings from our parallel investigations, ER stress and mitochondrial dysfunction emerge as key contributors to the disrupted Kv1.3 trafficking and retention in intracellular compartments. ODE exposure induced significant stress responses, including increased expression of spliced XBP1, ATF4, and GRP94, consistent with the unfolded protein response (UPR) activation observed under cellular stress (He et al., 2024; Walter & Ron, 2011). These findings suggest that ER stress as observed by us (Massey et al., 2021a) disrupts Kv1.3 protein trafficking, preventing its proper localization to the plasma membrane and likely trapping it within the ER and potentially other intracellular compartments. At the mitochondrial level, ODE exposure caused significant mitochondrial damage, characterized by mitochondrial hypertrophy, cristolysis, increased calcium sequestration bodies, and elevated mitochondrial stress markers such as MFN1, MFN2, and PINK1. These structural and functional impairments compromise mitochondrial bioenergetics, as evidenced by decreased mean oxygen consumption rate (OCR), basal respiration, and ATP production in Seahorse assays (Ratano et al., 2023). Given the known role of mitochondrial Kv1.3 (mitoKv1.3) in regulating mitochondrial membrane potential, ROS production, and apoptotic pathways, dysfunction at this level could result in Kv1.3 retention within the mitochondria, amplifying ROS production and pro-inflammatory signaling cascades (Capera et al., 2022; Wang et al., 2022).

Interestingly, despite the absence of enhanced surface Kv1.3 currents, we observed a significant increase in pro-inflammatory cytokine release, elevated NOX2 protein expression, and nitrite production. These outcomes are consistent with the hypothesis that intracellular Kv1.3 either in the ER or mitochondria—remains functionally active and contributes to these inflammatory pathways (Sarkar et al., 2020). Specifically, ER-localized Kv1.3 (ERKv1.3) may alter calcium signaling and protein folding homeostasis, exacerbating ER stress responses and downstream cytokine release (Pontisso et al., 2024). Mitochondrial Kv1.3 (mitoKv1.3) may enhance ROS production and mitochondrial dysfunction, contributing to NOX2 activation and sustained inflammatory signaling (Checchetto et al., 2019; Fukai & Ushio-Fukai, 2020). Together, these findings suggest a model where Kv1.3 is functionally active intracellularly but not on the plasma membrane, and its compartment-specific roles in mitochondria and ER drive the observed inflammatory and metabolic dysregulation. Inhibition of ER stress pathways and restoration of mitochondrial function could thus represent promising therapeutic avenues, not only to address Kv1.3 trafficking defects but also to mitigate the broader inflammatory and oxidative stress responses triggered by ODE exposure. This integrated perspective highlights the intricate interplay between Kv1.3 localization, ER stress, mitochondrial dysfunction, and inflammation as key drivers of microglial pathology in response to toxic insults.

The complexity of ODE, a heterogeneous mixture of organic and chemical components, may further contribute to the suppression of Kv1.3 surface expression. It is plausible that ODE triggers additional signaling pathways such as Fyn kinase, NF-κB activation, and downstream transcriptional effects that interfere with Kv1.3 trafficking or directly regulate its expression dynamics. Given this temporal complexity, future studies should evaluate Kv1.3 currents at multiple time-points post-ODE exposure and correlate intracellular accumulation with surface expression dynamics over time. From a therapeutic standpoint, pharmacological inhibition of Kv1.3 using PAP-1 has shown efficacy in reducing neuroinflammatory markers; however, its impact on Kv1.3 trafficking remains unclear. Investigating whether Kv1.3 inhibitors can restore proper protein trafficking and exploring combination therapies targeting Kv1.3 and oxidative stress pathways could offer novel therapeutic strategies. To gain deeper mechanistic insights, complementary techniques such as super-resolution microscopy or live-cell imaging should be employed to visualize Kv1.3 trafficking in real-time. Additionally, biotinylation assays can quantify surface versus intracellular Kv1.3 pools, and co-immunoprecipitation experiments may identify interacting partners influencing Kv1.3 localization and function. These findings emphasize the importance of investigating not only the localization but also the temporal regulation of Kv1.3 under conditions of ODE-induced neuroinflammation.

Concluding, this study advances our understanding of the multifaceted role of Kv1.3 in ODE-induced neuroinflammation. While intracellular Kv1.3 upregulation and its contribution to oxidative stress and cytokine secretion are evident, the lack of surface expression and associated Kv1.3 currents following ODE exposure suggest nuanced regulatory mechanisms that require further investigation. The findings underscore the potential of Kv1.3 as a therapeutic target, with pharmacological inhibition showing promise in mitigating key aspects of neuroinflammation, including nitrite (RNS marker) and cytokine release. Future directions should include a deeper investigation into Kv1.3 trafficking dynamics, the temporal progression of oxidative and inflammatory responses, and the role of other intersecting pathways in microglial activation. These insights will be crucial for developing targeted therapies to address neuroinflammation and its downstream effects in neurodegenerative diseases.

## Conclusions

Our study findings emphasize a central role of Kv1.3 in neuroinflammatory processes due to exposure to ODE. Although Kv1.3 expression increases intracellularly, its surface expression and functional currents remain unchanged due to protein trafficking issues linked to ER and mitochondrial damage. Kv1.3 amplifies oxidative stress and inflammatory responses through key pathways, such as NOX2-mediated nitrite production and cytokine release. Pharmacological inhibition of Kv1.3 with PAP-1 reduced oxidative stress markers and pro-inflammatory cytokine secretion, suggesting its potential as a therapeutic target. However, PAP-1’s partial efficacy indicates the involvement of additional signaling mechanisms, necessitating further research into complementary pathways. This study highlights Kv1.3 as a promising target for therapeutic intervention in ODE-induced neuroinflammation and future neurodegenerative disease research (Fig. 12).

**Figure 12.**
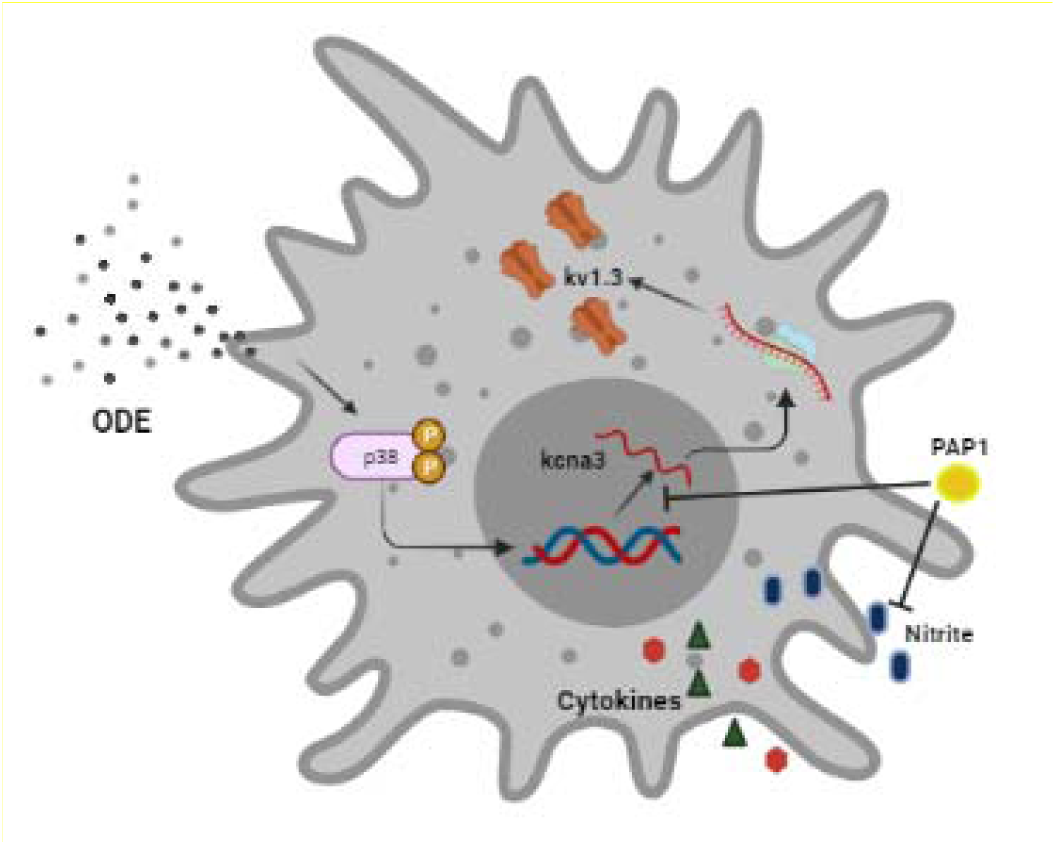
An overview of ODE exposure-induced kcna3 upregulation in primary microglia. ODE exposure mediated an increase in p38 MAPK phosphorylation, leading to upregulated *kcna3* expression. This was accompanied by elevated Kv1.3 protein levels in microglia following ODE stimulation. Additionally, ODE exposure triggered the secretion of pro-inflammatory cytokines and nitrite. Enhanced cellular capacitance was observed upon ODE stimulation, indicating an increase in microglial cell size associated with activation. Treatment with PAP-1 effectively reduced Kv1.3 protein levels, p38 MAPK phosphorylation, and nitrite secretion. However, PAP-1 treatment did not significantly alter pro-inflammatory cytokine secretion in response to ODE.

## Acknowledgment

We thank Dr T. Thippeswamy’s laboratory (Biomedical Sciences) for providing access to facilities. We also thank the Department of Biomedical Sciences for providing access to the core laboratory.

## Conflict of Interest

The authors declare that the research was conducted in the absence of any commercial or financial relationships that could be construed as a potential conflict of interest.

## Author Contributions

CC secured funding for the project, designed experiments, verified the data, offered intellectual input. NM conducted most of the experiments, wrote the first draft of the manuscript, and compiled the data. All other authors cross-verified the data, proof-read the manuscript with a significant contribution from the NM. NM, cross-verified and graphed the data.

## Funding

C.C. laboratory was funded through a startup grant through Iowa State University’s College of Veterinary Medicine and a pilot grant (5 U54 OH007548) from the CDC-NIOSH (Centers for Disease Control and Prevention-The National Institute for Occupational Safety and Health). A.G.K. laboratory is supported by the National Institutes of Health grants (ES026892, ES027245 and NS100090).

## Notes

### Competing Interest Statement

The authors have declared no competing interest.

