## Supplemental Data for "Exploring the Role of Kv1.3 and MAPK14 in Mediating Microglial Oxidative Stress and Neuroinflammation Following Organic Dust Exposure"

**Supplementary data**

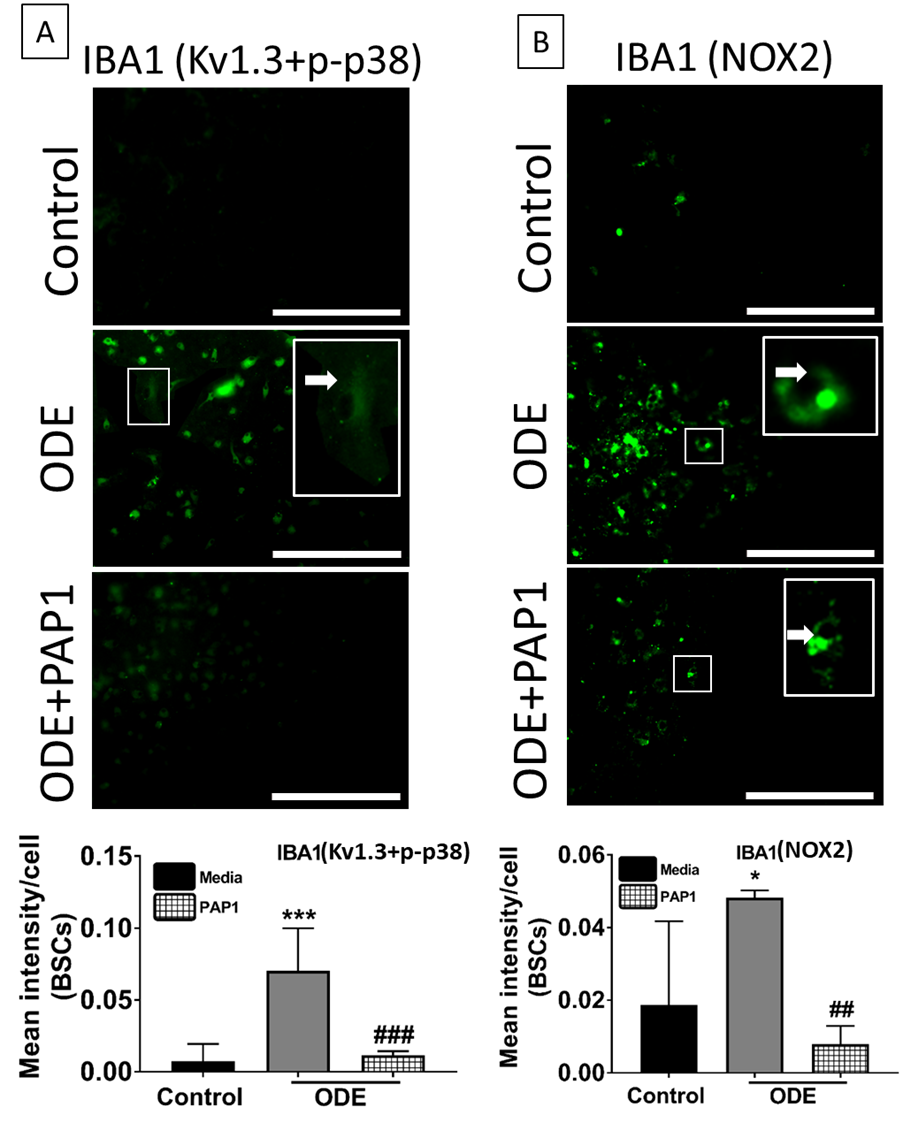

**Supplementary Figure 1 ODE induces IBA1 expression in BSCs.**

Treated (Table 13.1) BSCs were fixed with 4% paraformaldehyde. Following fixing, BSCs were stained with anti IBA1 (Fitc, green) antibody together with Kv1.3 and p-p38 (A) or NOX2 (B) antibodies. IBA1 expression in IHC was quantified, compared to control, ODE-exposed mice showed higher increased IBA1 staining intensity. PAP1 treatment significantly reduced IBA1 expression (n=3, * exposure effect, # PAP1 treatment effect, p ≤ 0.05, micrometer bar = 50 µm).

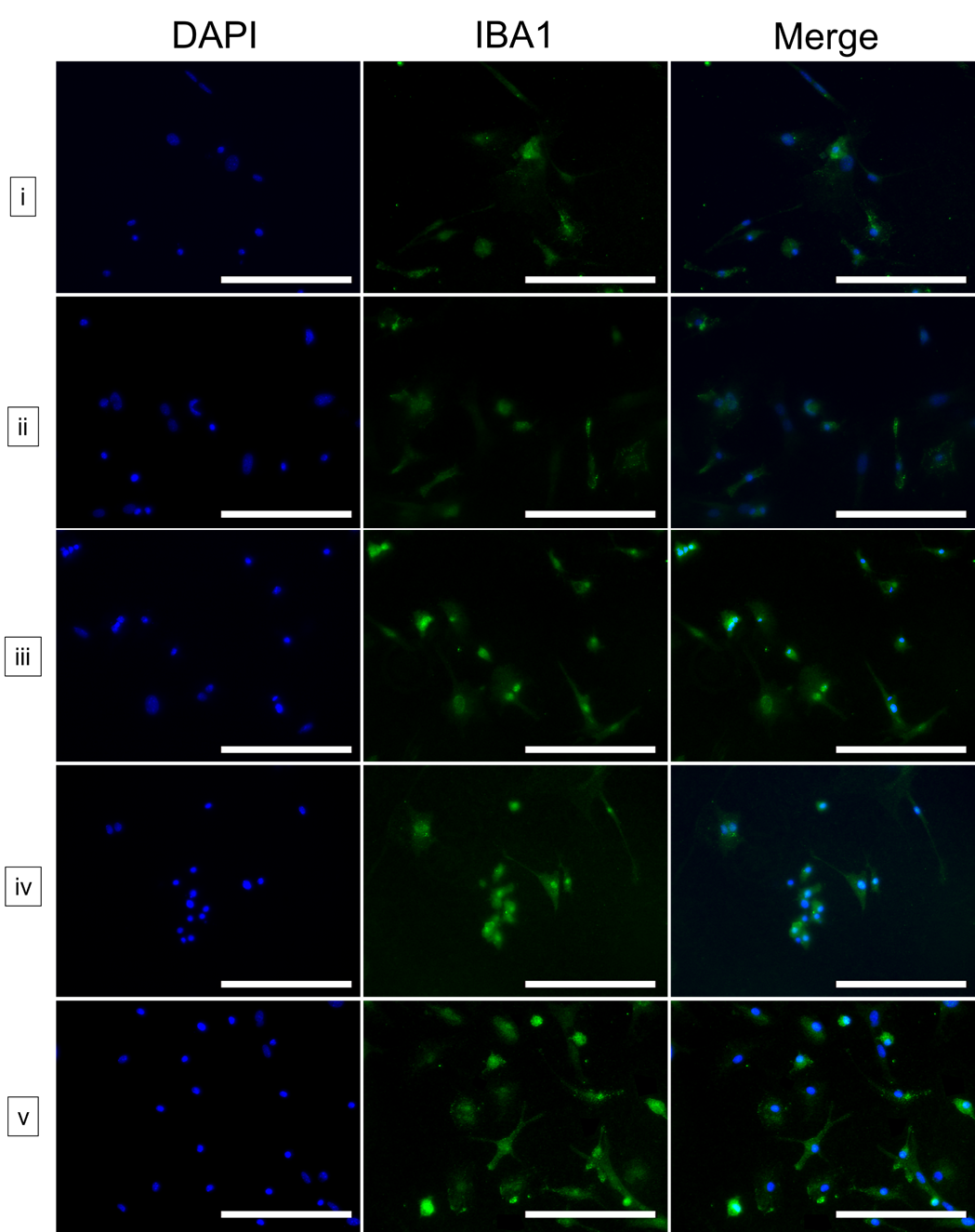

**Supplementary Figure 2. IBA1 positive cells following magnetic isolation**.

Following magnetic isolation of microglia from mixed glial culture, magnetically isolated cells were stained with anti-IBA1 (FITC) staining and counted. More than 96% of cells per field expressed IBA1, indicating a highly purified fraction.

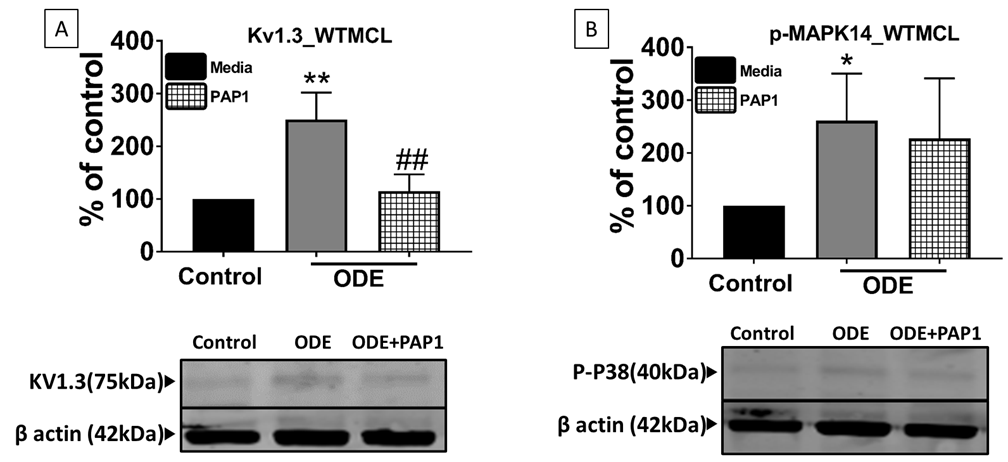

**Supplementary Figure 3. ODE induces Kv1.3 and p-MAPK14 protein expression in WTMCL**.

Whole cell lysate from WTMCL were processed for western blot analysis. KV 1.3, p-p38 MAPK and β-actin antibodies (house-keeping protein) detected 75 kDa, 40kDa and 42 kDa bands, respectively. Densitometry of normalized bands showed that, compared to controls, ODE exposure increased the Kv1.3 (A) and p-p38 MAPK (B) protein levels. PAP1 treatment significantly reduced both Kv1.3 (A) and p-p38 MAPK (B) protein level (n=3, * exposure effect, # PAP1 treatment effect, p ≤ 0.05).

**Supplementary Table 1. DEGs (Control Vs ODE)**

| Gene symbol | Gene name | Fold change | -log10(p value) |
| --- | --- | --- | --- |
| Ddx3y | DEAD (Asp-Glu-Ala-Asp) box polypeptide 3, Y-linked | 9.28778 | 9.6392 |
| Kdm5d | lysine (K)-specific demethylase 5D | 8.37495 | 9.6675 |
| Eif2s3y | eukaryotic translation initiation factor 2, subunit 3, structural gene Y-linked | 7.56009 | 7.99237 |
| Uty | ubiquitously transcribed tetratricopeptide repeat gene, Y chromosome | 6.4943 | 7.3404 |
| Mir3960 | microRNA 3960 | 3.0608 | 2.97837 |
| Kcna3 | potassium voltage-gated channel, shaker-related subfamily, member 3 | 2.97393 | 3.2481719 |
| Pth2 | parathyroid hormone 2 | 2.92944 | 2.72195 |
| Xylt1 | xylosyltransferase 1 | 2.84987 | 2.078611 |
| E2f2 | E2F transcription factor 2 | 2.72761 | 2.512473 |
| Abcc6 | ATP-binding cassette, sub-family C (CFTR/MRP), member 6 | 2.70606 | 2.02487 |
| Ly6g | lymphocyte antigen 6 complex, locus G | 2.55231 | 2.04767 |
| Hif1a | hypoxia-inducible factor 1, alpha subunit inhibitor | 2.53759 | 2.0941934 |
| Gm6878 | predicted gene 6878 | 2.50622 | 2.62555 |
| Rab44 | RAB44, member RAS oncogene family | 2.23999 | 1.47816 |
| Slpi | secretory leukocyte peptidase inhibitor | 2.19406 | 2.79084 |
| Gm5237 | predicted gene 5237 | 2.13247 | 1.92013 |
| Tstd1 | thiosulfate sulfurtransferase (rhodanese)-like domain containing 1 | 2.06113 | 3.0008 |
| Erdr1 | erythroid differentiation regulator 1 | -2.62243 | 2.57297 |
| Xist | inactive X specific transcripts | -11.0481 | 11.0693 |

**Supplementary Table 2. Gene ontology analysis of DEGs (Control-ODE)**

| Enriched pathway | Fold change | Category |
| --- | --- | --- |
| positive regulation of transcription from RNA polymerase II promoter | 4.2 | BP |
| inflammatory response | 3.3 | BP |
| positive regulation of transcription from RNA polymerase II promoter | 3.2 | BP |
| positive regulation of apoptotic process | 2.5 | BP |
| negative regulation of transcription from RNA polymerase II promoter | 2.5 | BP |
| oxidation-reduction process | 2.4 | BP |
| signal transduction | 2.1 | BP |
| signal transducer activity | 2.3 | MF |
| oxidoreductase activity | 2 | MF |

**Supplementary Table 3. Gene symbols for qRT-PCR validation**

forkhead box D1(Foxd1)

hypoxia inducible factor 1, alpha subunit(Hif1α)

interleukin 1 beta(Il1β)

nucleotide-binding oligomerization domain containing 2(Nod2)

transformation related protein 73(Trp73)

cytochrome b-245, beta polypeptide(Cybb)

adenosine deaminase(Ada)

ArfGAP with coiled-coil, ankyrin repeat and PH domains 1(Acap1)

cadherin 9(Cdh9)

G protein-coupled receptor 132(Gpr132)

RAB44, member RAS oncogene family (Rab44)

DEAD (Asp-Glu-Ala-Asp) box polypeptide 3, Y-linked (Ddx3y)

thbsomatostatin receptor 2(Sstr2)

keratin 2(Krt2)

E2F transcription factor 2(E2f2)

chemokine (C-X-C motif) ligand 2(Cxcl2)

secretory leukocyte peptidase inhibitor(Slpi)

thrombospondin 1(thbs1)

Proteoglycan 2 (PRG2)

Parathyroid hormone 1 receptor (PTHr)

xylosyltransferase 1(Xylt1)

potassium voltage-gated channel, shaker-related subfamily, member 3(Kcna3)

neuronal pentraxin receptor(Nptxr)

matrix metallopeptidase 25(Mmp25)

kinase suppressor of ras 2(Ksr2)

disrupted in schizophrenia 1(Disc1)

a disintegrin and metallopeptidase domain 8(Adam8)

CD300 molecule (Cd300)

erythroid differentiation regulator 1(Erdr1)

erb-b2 receptor tyrosine kinase 4(Erbb4)

nuclear factor, erythroid derived 2(Nfe2/NRF2)

NK-3 transcription factor, locus 1 (Drosophila)(Nkx3-1)

cytochrome c oxidase subunit I(COX1)

NADH dehydrogenase subunit 2(ND2)

NADH dehydrogenase subunit 1(ND1)

plasminogen activator, urokinase receptor(Plaur)

neuronal pentraxin receptor(Nptxr)

matrix metallopeptidase 9(Mmp9)

bone morphogenetic protein receptor, type II (serine/threonine kinase)(Bmpr2)

**Supplementary Table 4. Details of statistical analysis applied for each experiment/figure.**

| Figure | Panel | Test |
| --- | --- | --- |
| 6.1 | A-B | one-way ANOVA; Tukey’s post-hoc, n=3 mice |
| 6.2 |  | one-way ANOVA; Tukey’s post-hoc, n=3 mice |
| 6.3 |  | one-way ANOVA; Tukey’s post-hoc, n=3 mice |
| 6.4 | A-C | one-way ANOVA; Tukey’s post-hoc, n=3 mice |
| 6.5 | A&D | two-way ANOVA; Tukey’s post-hoc; n=3 wells (4-5 slices/ well) (6 well plate) |
|  | B&E | two-way ANOVA; Tukey’s post-hoc; n=3 wells (24 well plate) |
|  | C&F | two-way ANOVA; Tukey’s post-hoc; n=3 wells (24 well plate) |
| 6.6 | B | two-way ANOVA; Tukey’s post-hoc; n=3 wells (4-5 slices/ well) (6 well plate) |
|  | D&F | two-way ANOVA; Tukey’s post-hoc; n=3 wells (24 well plate) |
| 6.7 | B | two-way ANOVA; Tukey’s post-hoc; n=3 wells (4-5 slices/ well) (6 well plate) |
|  | D | two-way ANOVA; Tukey’s post-hoc; n=3 wells (24 well plate) |
| 6.8 | B | two-way ANOVA; Tukey’s post-hoc; n=3 wells (4-5 slices/ well) (6 well plate) |
|  | D&F | two-way ANOVA; Tukey’s post-hoc; n=3 wells (24 well plate) |
| 6.9 | A | two-way ANOVA; Tukey’s post-hoc; n=3 wells (4-5 slices/ well) (6 well plate) |
|  | B-C | two-way ANOVA; Tukey’s post-hoc; n=3 wells (24 well plate) |
| 6.10 | B,C,D | two-way ANOVA; Tukey’s post-hoc; n=3 wells (24 well plate) |
| 6.11 | A-B | two-way ANOVA; Tukey’s post-hoc; n=3 wells (4-5 slices/ well) (6 well plate) |
| 6.12 | A-B | two-way ANOVA; Tukey’s post-hoc; n=3 wells (24 well plate) |
